# Deep Learning Structural Ensembles as Proxies for Protein Flexibility

**DOI:** 10.64898/2026.05.16.725658

**Authors:** Mehmet Tahir Tunc, Ayten Dizkirici Tekpinar, Mustafa Tekpinar

## Abstract

Protein dynamics are essential to biological function, yet understanding whether deep learning models contain information about these dynamics remains an open question. In this study, we quantitatively investigate the capacity of deep learning structure generation methods to predict protein flexibilities by directly comparing residue-level mean squared fluctuation (MSF) profiles derived from structural ensembles with experimental or simulation-informed flexibility profiles. We assembled four diverse benchmark datasets representing different types of structural information, including 70 NMR ensembles, 43 X-ray crystallographic protein pairs in two distinct conformational states, 82 high-resolution cryo-EM structures, and molecular dynamics simulations of 10 proteins. Utilizing AlphaFold3, AlphaFold2, and RosettaFold to generate multiple structural models, we applied ranksort normalization to place the profiles on a comparable scale and quantified similarity primarily using cosine and Pearson similarities. Our results demonstrate that the flexibility predictions from deep learning-generated models agree well with experimental data, suggesting that fluctuations in these predicted ensembles can serve as effective proxies for protein flexibility. Notably, AlphaFold3 consistently produced the best results across the datasets. We also observed that flexibility prediction accuracy generally improves as the number of models increases up to 15, and our findings remain robust even when terminal residues are excluded from the analysis. To facilitate broader application, we provide three publicly accessible Jupyter Notebooks to calculate MSF from deep learning outputs. Ultimately, this work provides evidence that deep learning structural ensembles can serve as proxies for protein flexibility.

## INTRODUCTION

Deep learning-based protein structure prediction has transformed structural biology and its allied disciplines^1–4^. The number of accessible protein structures has increased by almost three orders of magnitude, from about 200000 to hundreds of millions in the last few years. The abundance of the protein structures expanded our understanding of proteins in all structure related fields. However, understanding if these models contain information about protein dynamics remains an open question.

Proteins are dynamic macromolecules and their dynamics have strong implications in their functioning. Determining different conformational states of proteins, their transitions between them and flexible regions in proteins facilitating these conformational transitions are some major research areas in protein dynamics. Accordingly, protein dynamics has been investigated extensively using both experimental and computational approaches^5–11^.

Due to the large success of deep learning in protein structure prediction, predicting protein dynamics using deep learning has become a major research focus in recent years. Protein dynamics can be broadly categorized into two partially overlapping areas: protein conformational sampling and protein flexibility investigations. Initial studies on protein dynamics and deep learning protein structure prediction algorithms focused on obtaining different conformational states of proteins by modifying multiple sequence alignment (MSA) files^12–15^. However, the success rates of these approaches were limited. Understanding and predicting protein flexibilities represented an additional frontier in the field of protein dynamics. The potential relation between protein flexibilities and pLDDT have also been investigated in many studies ^16–18^. However, many of protein flexibility research rely on indirect indicators, such as confidence scores etc., rather than comparing ensemble-derived flexibility measures directly.

In principle, protein flexibility can be directly measured from an ensemble of structures using a widely used metric called mean squared fluctuations (MSF). Experimental methods that can generate multiple conformational states of a protein can be used to compute the MSF. Furthermore, even though normal modes are small fluctuations around an equilibrium, they can also constitute an ensemble for the MSF calculations. Molecular dynamics simulations can provide large ensembles for the MSF calculations. The source of the structural ensemble can also be models from deep learning structure prediction methods like AlphaFold3, AlphaFold2 or RosettaFold. If the MSF from the experiments/simulations and deep learning structure prediction algorithms are compared, this may allow us to quantify the extent of (dis)agreement between these resources. As a result, the capacity of deep learning structure generation methods to predict protein flexibilities can be investigated quantitatively in this way.

In this study, we address this question by directly comparing residue-level MSF profiles derived from deep learning structural ensembles with experiment or simulation-derived flexibility profiles. We assembled four benchmark datasets based on NMR, X-ray crystallography, cryo-EM structures and molecular dynamics (MD) simulation ensembles. Since the available experimental information differs across methods, our benchmarks include both direct ensemble-based measures and structure-derived proxies of flexibility. After computing MSF profiles from experiments, MD simulations and deep learning-generated structures, we applied ranksort normalization to place the profiles on a comparable relative scale and quantified similarity primarily using cosine and Pearson similarities. We investigated impact of model numbers and terminal effects to quantify robustness of our approach. This framework allows us to test whether current deep learning structural models can be used as proxies of protein flexibility across diverse proteins and experimental/simulation contexts.

## MATERIALS AND METHODS

### Materials

#### Dataset overview

We compiled four benchmark datasets representing different types of experimentally or simulation derived structural information: 70 NMR ensembles, 43 X-ray crystallographic protein pairs solved in two conformational states, 82 high-resolution cryo-EM structures and MD simulations of 10 proteins, each one with three replicas. These datasets differ in the degree to which they provide direct ensemble information; therefore, they should be interpreted as complementary rather than identical measures of protein flexibility.

#### NMR Dataset

We used an NMR dataset that has been used in a previous study^19^. We selected 70 proteins of various sizes (Supporting Information Table S1) from that dataset. The smallest protein (PDB ID: 6svc^20^) has 35 amino acids and the largest protein (PDB ID: 2lf2^21^) contains 175 amino acids. Only 3 proteins (PDB IDs: 1yez^22^, 1pqx and 2m7u^23^) contain 10 NMR models, while the remainder contain 20 models. The dataset is quite diverse in terms of the sequence identities (**Figure S1** A).

#### X-Ray Dataset

The X-Ray dataset contains 43 proteins that have been obtained in two conformational states (Table S2). These proteins have been widely used in previous studies^24–26^. The smallest protein (PDB ID1: 1e7xA- PDB ID1:1dzsB) in the dataset has 129 amino acids, while the biggest protein (PDB ID1: 3kqnA- PDB ID1: 8ohmA) has 435 residues. Protein pairs with more than 500 residues were excluded because RosettaFold server cannot generate models for proteins beyond that threshold. Root Mean Squared Distance (RMSD) between protein conformations varies from 0.29 Å to 16.08 Å. This dataset is also quite diverse in terms of the sequences (**Figure S1** B).

#### Cryo-EM Dataset

This dataset contains 82 proteins, whose details given in Table S3. We selected cryo-EM structures with≤3 Å resolution using advanced search in rcsb.org. Polymer entity type was set to protein and number of assemblies was set to 1 in the advanced search. Only proteins with less than 500 residues were selected due to the aforementioned limitation in the RosettaFold server. Sequence diversity of the dataset is provided in (**Figure S1** C).

#### Molecular Dynamics (MD) Simulations Dataset

We selected 10 proteins for MD simulations dataset. Five of these proteins were the simulations investigated by one of the authors previously^27–29^. The simulations of the other five protein were obtained from the ATLAS database^30^, which is a standardized all-atom MD simulations database at https://www.dsimb.inserm.fr/ATLAS. Each protein has three replicas of the simulations, presented as simulation 1, 2 and 3 in the paper. The size of the proteins in our dataset ranges from 71 amino acids to 453 amino acids. Essential details of the proteins and the simulations are given in Table S4, while sequence identities are provided in **Figure S1** D.

#### Pre-processing, residue mapping, and structural superposition

For each protein, experimental structures and predicted models were mapped to a common 0-based residue index using the deposited PDB/mmCIF and the input FASTA sequence because residue IDs in the pdb files and deep learning models may not always match. Prior to the MSF calculations, all models within each ensemble (experimental, simulated or predicted) were rigid-body superimposed using a least-squares Calpha alignment to remove global translation/rotation components.

### Methods

#### Calculation of Deep Learning Predicted Structural Models

We used three deep learning structure prediction algorithms to generate multiple models for each protein. Twenty AlphaFold3 structural models were generated using the weights provided Google Deepmind^31^. We used alphafold3_tools to convert fasta files to json format for AlphaFold3^32^. modelSeeds: [1, 2, 3, 4] parameter was used in the json file to generate 20 models. Similarly, twenty AlphaFold2 structural models were obtained from ColabFold (v1.6.1) implementation at https://colab.research.google.com/github/sokrypton/ColabFold/blob/main/AlphaFold2.ipynb^33, 34^ using ‘num_seeds=4’ and ‘use_dropout=True’ parameters. However, we used the first five models with the highest ranks as standard inputs both in AlphaFold3 and AlphaFold2 because the servers provide five models by default. Since one cannot generate more than five models with RosettaFold server, only five models were obtained from robetta server at https://robetta.bakerlab.org/^35^.

#### Mean Squared Fluctuations (MSF)

MSF or its square root (RMSF) are commonly used metrics to quantify protein flexibility^36–41^. MSF were calculated using the following equation:

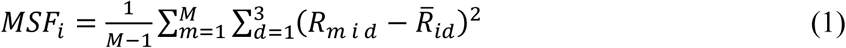

In this equation, 𝑀𝑆𝐹_𝑖_ is mean squared fluctuation of residue *i*. 𝑀 is number of models in the ensemble, 𝑅_𝑚𝑖_ is 3D coordinate vector of residue *i* in model m and 𝑅̅_𝑖𝑑_ is mean coordinate of residue *i* in dimension *d*. We used only coordinates of Calpha atoms for the MSF calculations. The models may come from experimental structures, MD simulations or deep learning structure prediction algorithms like AlphaFold3, AlphaFold2 or RosettaFold. We should note that 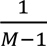 factor in the equation becomes unimportant when a normalization is applied to the MSF values.

#### Normalization of the Mean Squared Fluctuations

It is necessary to normalize the MSF to compare MSF from different sources. There are several normalization methods, such as minmax or ranksort normalization, that can be applied for this purpose. Ranksort normalization was preferred over minmax normalization because it is robust to outliers arising from highly flexible terminal residues or disordered loops, which can dominate minmax scaling and obscure the overall flexibility pattern in the protein. In ranksort normalization, MSF are ranked according to their values. The ranked values are divided by the length of the array to obtain values in a standard scale between [0, 1].

#### Coarse-Grained Normal Mode Ensemble Calculation with Anisotropic Network Model (ANM)

In ANM, all Calpha atoms of a protein within a cutoff radius (Rc=15 Å in this study) are connected with springs of uniform stiffness (γ) and they are assumed to interact through a simple Hookean potential given below^40^:

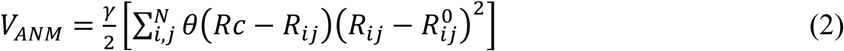

*R^0^_ij_* is equilibrium distance between Calpha atoms of residue *i* and *j*, while *R_ij_* is the distance between Calpha atoms of residue *i* and *j* at any moment. 𝜃(𝑅𝑐 − 𝑅_𝑖𝑗_) is Heaviside step function to include only Calpha atoms within the cutoff radius (Rc=15 Å). A hessian matrix can be constructed from this potential. The hessian matrix has to be diagonalized to find eigenvalues and eigenvectors to obtain normal modes. The eigenvectors corresponding to translations and rotations are eliminated and the remaining eigenvectors constitute the normal mode ensemble. The low frequency modes are generally collective motions in a protein. It has been shown that the normal modes obtained with this type of potential can be used to predict observed conformational changes of proteins as well as fluctuations of proteins around native conformations ^25, 40, 42^. All normal modes and MSF calculations from these modes were performed with Prody^43, 44^ and sedy (https://gitlab.com/tekpinar/sedy) packages. Finally, it is important to note that the normal modes are small fluctuations around an equilibrium position by definition^45^. Even though they do not sample different conformations of proteins, they are quite good at predicting protein flexibilities using these small fluctuations.

#### Quantifying Similarity of the MSF from Different Sources

Cosine similarity is a metric used to measure how similar two vectors are, regardless of their magnitude. It is formulated as follows for vectors A and B:

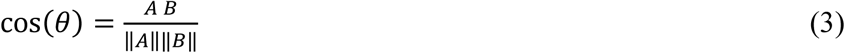

‖𝐴‖ and ‖𝐵‖ are magnitudes of vectors *A* and *B*. When the vectors point in the same direction (0° angle), cosine similarity is 1.0, which means perfect similarity. When the vectors point in opposite directions (180° angle), cosine similarity is −1.0. When the vectors are orthogonal (90° angle), the vectors are unrelated and cosine similarity is zero. We used cosine similarity to quantify similarity of MSF from experimental sources and MD simulations with three deep learning structure prediction methods. In this case, vector A represents our MSF from a reference method (experiment or simulation) and vector B represents the MSF from a deep learning ensemble. We should note that cosine similarity has been used to quantify similarity of MSF, RMSF or direction of conformational changes in previous studies as well^25, 46^. Furthermore, we used Pearson similarity, which measures linear correlations between two sets of data as a secondary metric to quantify the similarities. Like cosine similarity, Pearson similarity also ranges between [-1.0, 1.0]. While 1.0 value indicate a high correlation (or anti-correlation for negative values), zero denotes no correlation. It is important to note that Pearson similarity may be low when the correlations are non-linear.

## RESULTS

### MSF from the NMR Dataset and the Deep Learning Predicted Structural Models

We calculated MSF using the procedure explained in the Materials and Methods section. The MSF were ranksort normalized and added to the B-factors column of the PDB files. Then, the proteins were visualized with Pymol^47^ and colored according to the ranksorted MSF values. An example visualization for protein 2lah is provided in **Figure 1** A. Visual inspection demonstrates that one terminal region of the protein is quite flexible. This flexibility can be observed in AlphaFold3, AlphaFold2 and RosettaFold models as well. Furthermore, we plotted the experimental and computed flexibilities for 2lah as a 2D plot in **Figure 1** B. AlphaFold3 predicted flexibility has 0.988 cosine similarity to the experimental data while the AlphaFold2 has 0.911 cosine similarity. Cosine similarity of RosettaFold to the experimental data is 0.958. All of the flexibility predictions for this protein have quite good agreement with the experimental data. Finally, we computed cosine similarities of all proteins with the experimental data as bar plots in **Figure 1** C. Average cosine similarity of 70 proteins for AlphaFold3 is 0.940±0.034. The average value reduces to 0.915±0.041 for AlphaFold2 and 0.918±0.040 for RosettaFold (**Figure 2** A). Pearson similarity data is provided **Figure 2** B and **Figure S2.** Pearson similarity values are slightly reduced but they follow similar patters with cosine similarity data. Overall, the flexibility predictions from deep learning generated models agree well with the experimental data. These results demonstrate that fluctuations in deep learning predicted structural models can be used as proxies of protein flexibilities observed in the NMR ensembles. Furthermore, it is worthy of noting that AlphaFold3 models produce the best results in this dataset.

**Figure 1.**
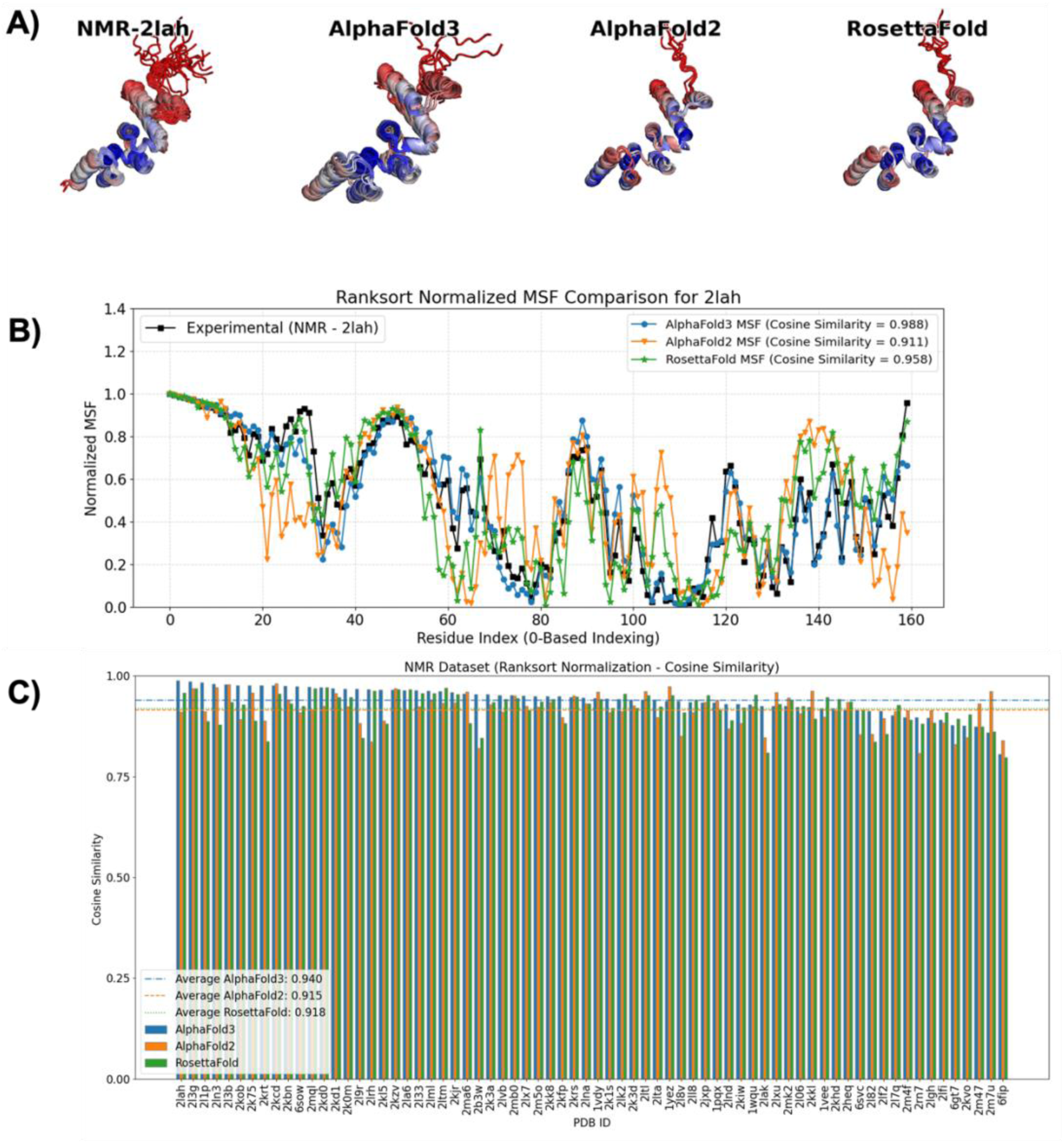
Ranksorted mean squared fluctuations (MSF) from NMR method and three deep learning structure prediction methods (AlphaFold3, AlphaFold2 and RosettaFold). A) Projections of ranksorted MSF onto protein structures for 2lah. Blue-White-Red color palette is used for the projections, where blue indicates low flexibility and red indicates high flexibility. B) 2D comparison of the experimental and the computed MSF for 2lah. Black line (with squares) is for the experimental data, blue line (with circles) is for AlphaFold3, orange line (with inverse triangles) is for AlphaFold2 and green line (with stars) is for RosettaFold. C) Cosine similarity of the experimental and the computed MSF for 70 proteins in the NMR dataset. AlphaFold3 bars are blue, AlphaFold2 bars are orange and RosettaFold bars are green. Averages of the cosine similarities over the entire dataset are also provided as horizontal lines for AlphaFold3 (blue dot-dashed line), AlphaFold2 (orange dashed line) and RosettaFold (green dotted line).

**Figure 2.**
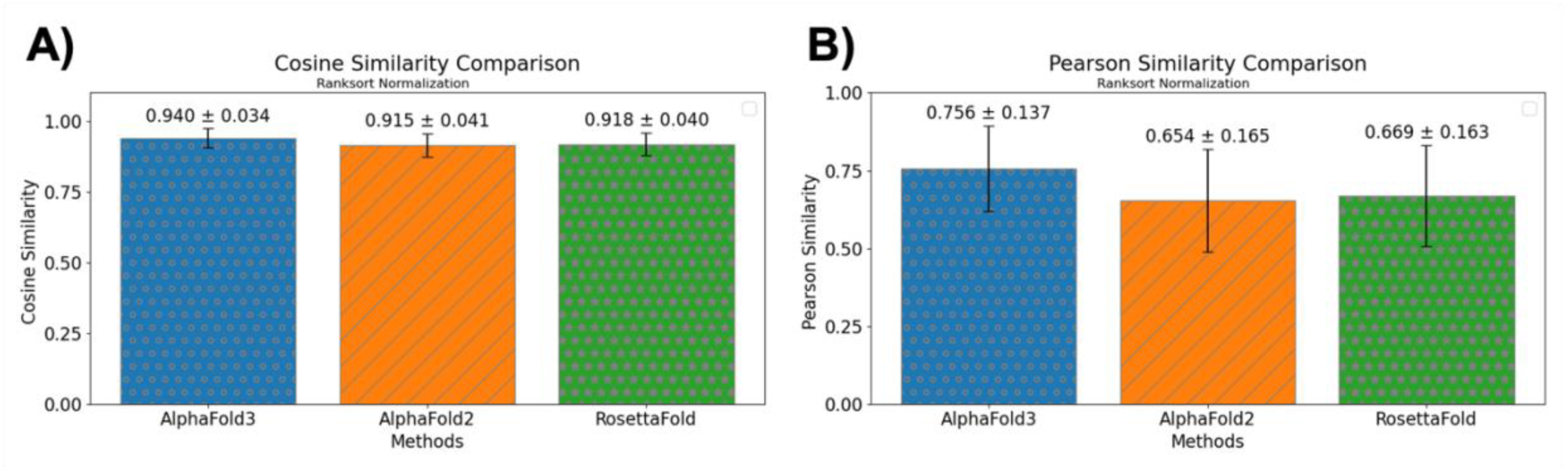
Average similarities with standard deviations as error bars for different approaches to obtain MSF from the NMR experimental data. Blue bars with gray circles are for AlphaFold3, orange bars with gray stripes are for AlphaFold2 and green bars with gray stars are for RosettaFold. A) Average cosine similarities B) Average Pearson similarities.

### Impact of Number of Deep Learning Structural Models on the MSF of the NMR Dataset

An important point here is to investigate impact of number of deep learning structural models on the MSF predictions. We calculated the MSF from 5, 10, 15 and 20 models, where the models are ordered according to their ranks, using AlphaFold3 and AlphaFold2. Then, we calculated average cosine and Pearson similarities. The average similarities increase in AlphaFold3 when more models are employed in the MSF calculations (**Figure 3** A left and right panels). On the other hand, the average similarities saturate after 15 model in AlphaFold2 calculations (**Figure 3** B left and right panels).

**Figure 3.**
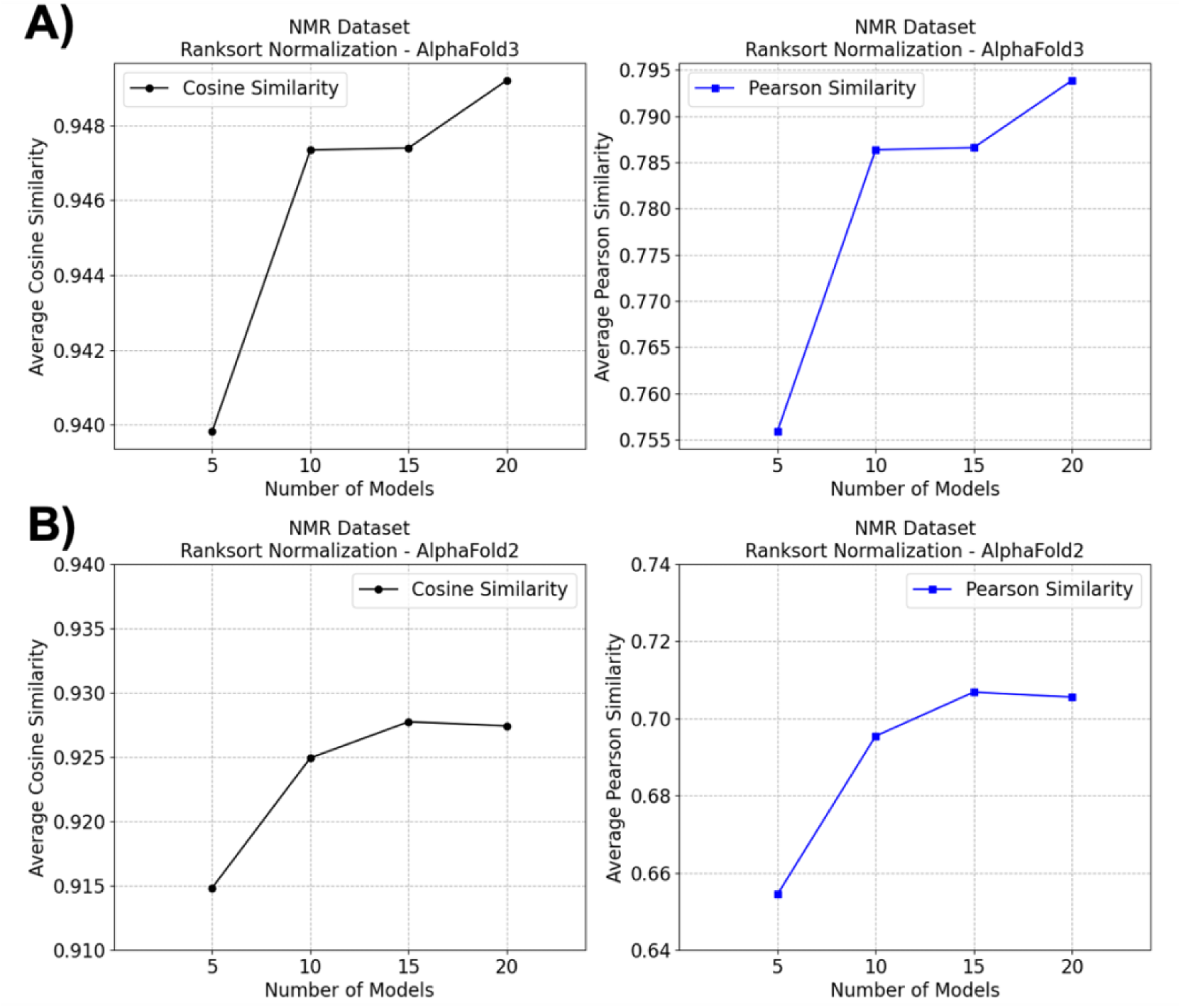
Impact of number of deep learning structural models on the NMR dataset, measured with cosine similarity (left panel) and Pearson similarity (right panel) for A) AlphaFold3 B) AlphaFold2.

### MSF from the X-Ray Dataset and the Deep Learning Predicted Structural Models

#### MSF from X-Ray Dual Conformations and Deep Learning Predicted Structural Models

Unlike proteins obtained with NMR method, we do not have many models for proteins obtained with X-ray crystallography. However, we have a set of proteins in two conformational states obtained with X-ray crystallography and the experimental MSF can be calculated from these dual conformational states. An example comparison between the experimental and deep learning-based MSF for catabolite gene activator protein (PDB IDs:3fwe^48^ chain B, 4hzf^49^ chain A) is provided in **Figure 4** A. Here, we can observe that some helices are highly flexible. Furthermore, AlphaFold3 flexibilities are in very good agreement with the experimental MSF (**Figure 4** B). A bar plot is provided to demonstrate cosine similarity of 43 proteins in the X-ray dataset (**Figure 4** C). Similar to the NMR dataset results, AlphaFold3 is the best method for predicting protein flexibilities with an average cosine similarity of 0.888±0.048. AlphaFold2 (with 0.871±0.051 cosine similarity) and RosettaFold (with 0.822±0.037 cosine similarity) follow the AlphaFold3 predictions. The Pearson similarities shown in **Figure S3** follow similar trends with the cosine similarities. To summarize, the x-ray dataset utilizing dual conformations of the proteins provide further evidence that protein flexibilities from deep learning generated ensembles are in decent agreement with the experimental data.

**Figure 4.**
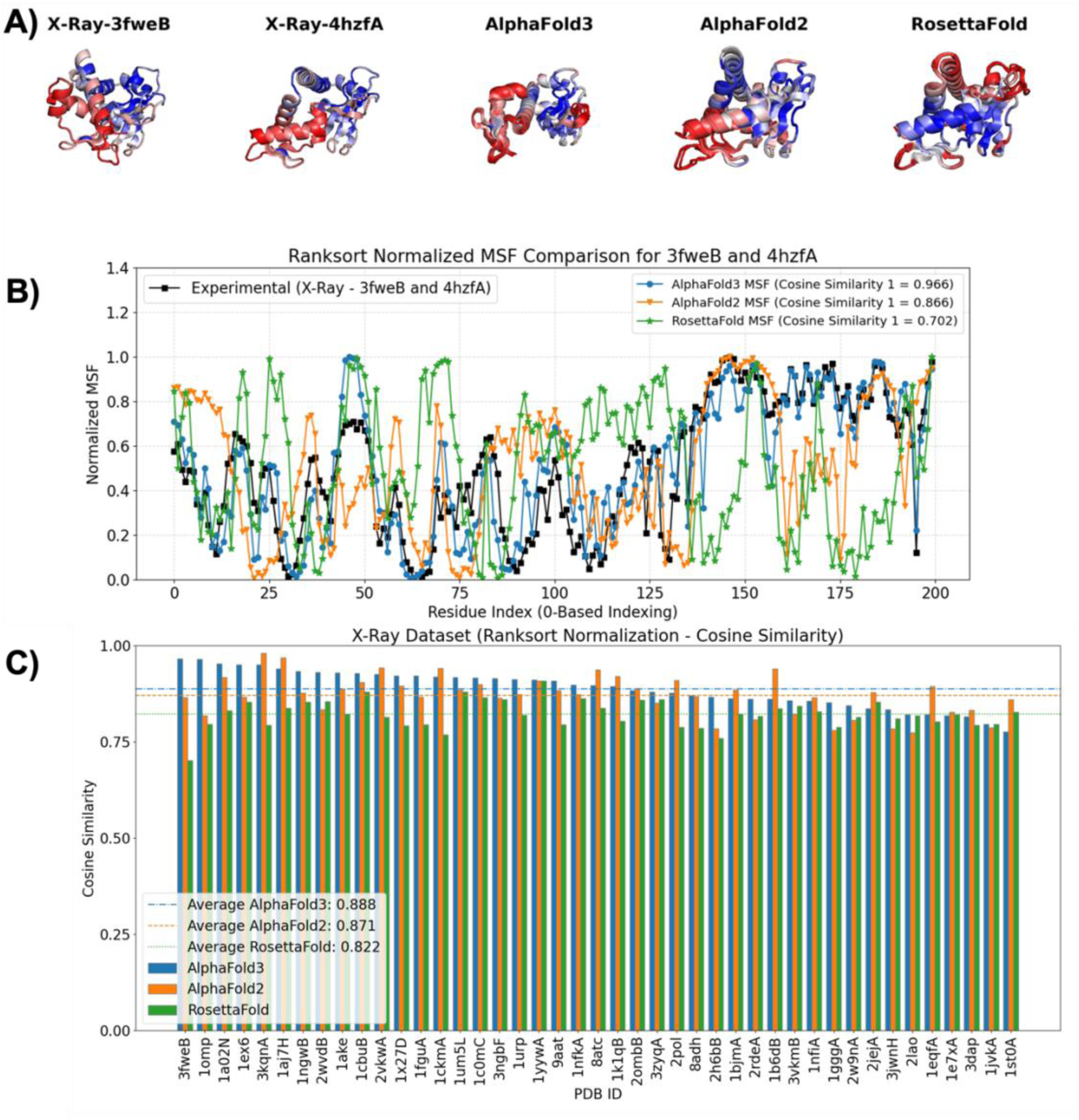
Ranksorted mean squared fluctuations (MSF) from dual conformations of X-Ray structures and three deep learning structure prediction methods (AlphaFold3, AlphaFold2 and RosettaFold). A) Projections of the ranksorted MSF onto protein structures for 3fweB and 4hzfA. Blue-White-Red color palette is used for the projections, where blue indicates low flexibility and red indicates high flexibility. B) 2D comparison of the experimental and the computed MSF for 3fweB and 4hzfA. Black line (with squares) is for the experimental data, blue line (with circles) is for AlphaFold3, orange line (with inverse triangles) is for AlphaFold2 and green line (with stars) is for RosettaFold. C) Cosine similarity of the experimental and the computed MSF for 43 proteins in the X-Ray dataset. AlphaFold3 bars are blue, AlphaFold2 bars are orange and RosettaFold bars are green. Averages of the cosine similarities over the entire dataset are also provided as horizontal lines for AlphaFold3 (blue dot-dashed line), AlphaFold2 (orange dashed line) and RosettaFold (green dotted line).

Due to the restriction of maximum 500 residues in RosettaFold server, we have not included any protein longer than this limit in the 43 proteins dataset. However, we wondered the cosine similarity between the experimental MSF and the deep learning structural ensemble MSF for longer proteins. Consequently, we identified two protein pairs obtained via X-ray crystallography: 1d6m^50^-1i7d^51^ with 603 amino acids and 1lfh^52^-1lfg^53^ with 691 amino acids. We calculated the experimental MSF curves from two conformations of the proteins. Then, we computed deep learning MSF from AlphaFold3 and AlphaFold2 models. The MSF from two sources were projected onto the protein structures and their 2D plots were created (**Figure S4**). Our results demonstrate that the MSF curves and their projections from the experimental MSF and the deep learning MSF agree with each other to a large extent. The cosine similarity of the curves is more than 0.806 for both of the protein pairs. To summarize, we observe a decent agreement for two proteins with more than 500 residues.

#### Experiment-Derived MSF with Other Approaches for the X-Ray Dataset

We used two conformations to determine protein flexibilities, which may not be optimal. As a result, we need methods that can generate conformational ensembles based on the experimental data. Coarse-grained normal mode analysis is a method that can generate 3N-6 normal modes, where N is number of its Calpha atoms. Historically, this approach has been used widely to study protein domain motions, conformational changes, MSF calculation and protein flexibilities. In this approach, each normal mode analysis needs only an input structure for the normal mode ensemble generation. After normal mode ensemble calculation, we calculated experimental-derived MSF using all normal modes for two conformations of the 43 proteins in the x-ray dataset (**Figure S5** for cosine similarities and **Figure S6** for Pearson similarities). The average cosine similarity increases to 0.910 (0.911) for AlphaFold3 when the first (second) conformations of the proteins are used in all normal mode calculations. Furthermore, we investigated whether low-frequency collective motions are the major contributors to protein flexibility. As a result, we recalculated protein flexibilities utilizing only the lowest frequency 10 normal modes (**Figure S7** for cosine similarities and **Figure S8** for Pearson similarities). The results of all modes and only 10 modes are quite similar, although 10 normal mode approach generates better MSF for this dataset. This point demonstrates that the MSF from the collective motions are embedded in the MSF of deep learning structural ensembles to a large extent.

In terms of averages of the similarities, MSF from normal mode-based approach generate a better agreement with deep-learning-based MSF (**Figure 5** A-B-C and **Figure S9** A-B-C). Previous studies demonstrated that normal modes, in particular the lowest frequency normal modes, show direction of different conformational transitions and domain motions^25, 54^. As a result, it is not surprising to see that normal-mode-based MSF produces results comparable (or better) protein flexibilities to the all normal mode approach.

**Figure 5.**
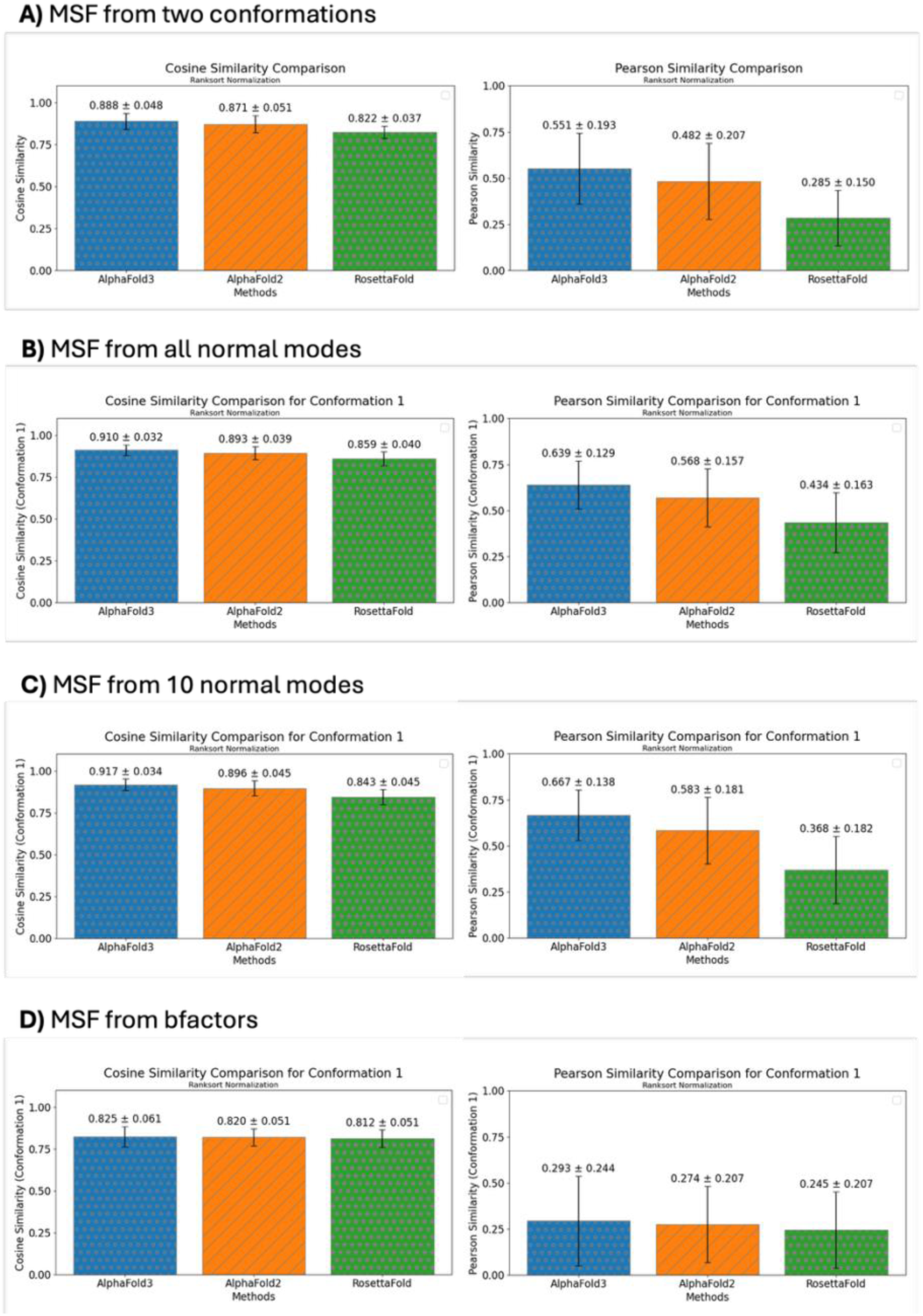
Average of similarities for different approaches to obtain MSF from the first conformation set of the X-Ray experimental data. The average cosine similarities are given in the left panel and the average Pearson similarities are provided in the right panel. The average was taken over the similarities of 43 proteins. Blue bars with gray circles are for AlphaFold3, orange bars with gray stripes are for AlphaFold2 and green bars with gray stars are for RosettaFold. A) Two experimental protein conformations were used for the MSF calculations. B) All normal modes of the first conformations was used for the MSF calculations. Only Calpha atoms were used for normal mode analysis. C) Only 10 lowest eigenvalue normal modes of each conformation were used for normal mode analysis. Only Calpha atoms were used for normal mode analysis. D) Bfactors were ranksort normalized and they were used as a proxy for the MSF.

One classical approach to extract protein flexibilities from experimental x-ray structures is to use their bfactors as an estimator of the experimental protein flexibility. Due to this reason, we extracted bfactors from X-ray structures and ranksort normalized them. Their average cosine and Pearson similarities with the MSF from deep learning predicted structural ensembles is given in **Figure 5** D and **Figure S9** D. The similarity is lower than the other approaches presented in **Figure 5** A, B and C (see **Figure S9** A, B and C for a comparison over the second conformations). These results support the idea that bfactors from crystallographic data may not be the most complete representatives of protein flexibilities as previously pointed out ^55, 56^.

#### Impact of Number of Deep Learning Structural Models on the MSF of the X-Ray Dataset

We calculated MSF from dual protein conformations, all normal modes, 10 lowest frequency modes and bfactors in the previous subsection for the proteins in the X-ray dataset. Our results demonstrated that the MSF from the normal mode approach agree well with the MSF from deep learning structural ensembles. We had used only five models with the highest rank as our deep learning structural ensemble. A systematic analysis of influence of number of deep learning structural models (5, 10, 15 and 20) on MSF similarities for AlphaFold3 and AlphaFold2 methods is presented in **Figure 6** for all normal mode approach. We observe that the average MSF similarity increases when up to 15 models are included. The average similarity starts to deteriorate (for AlphaFold3) or saturate (AlphaFold2) beyond this point. The results are quite similar when 10 normal modes are employed for the MSF calculations (**Figure S10)**. The impact results from dual conformational states and bfactor-based MSF are provided in **Figure S11** and **Figure S12** for further inquiry.

**Figure 6.**
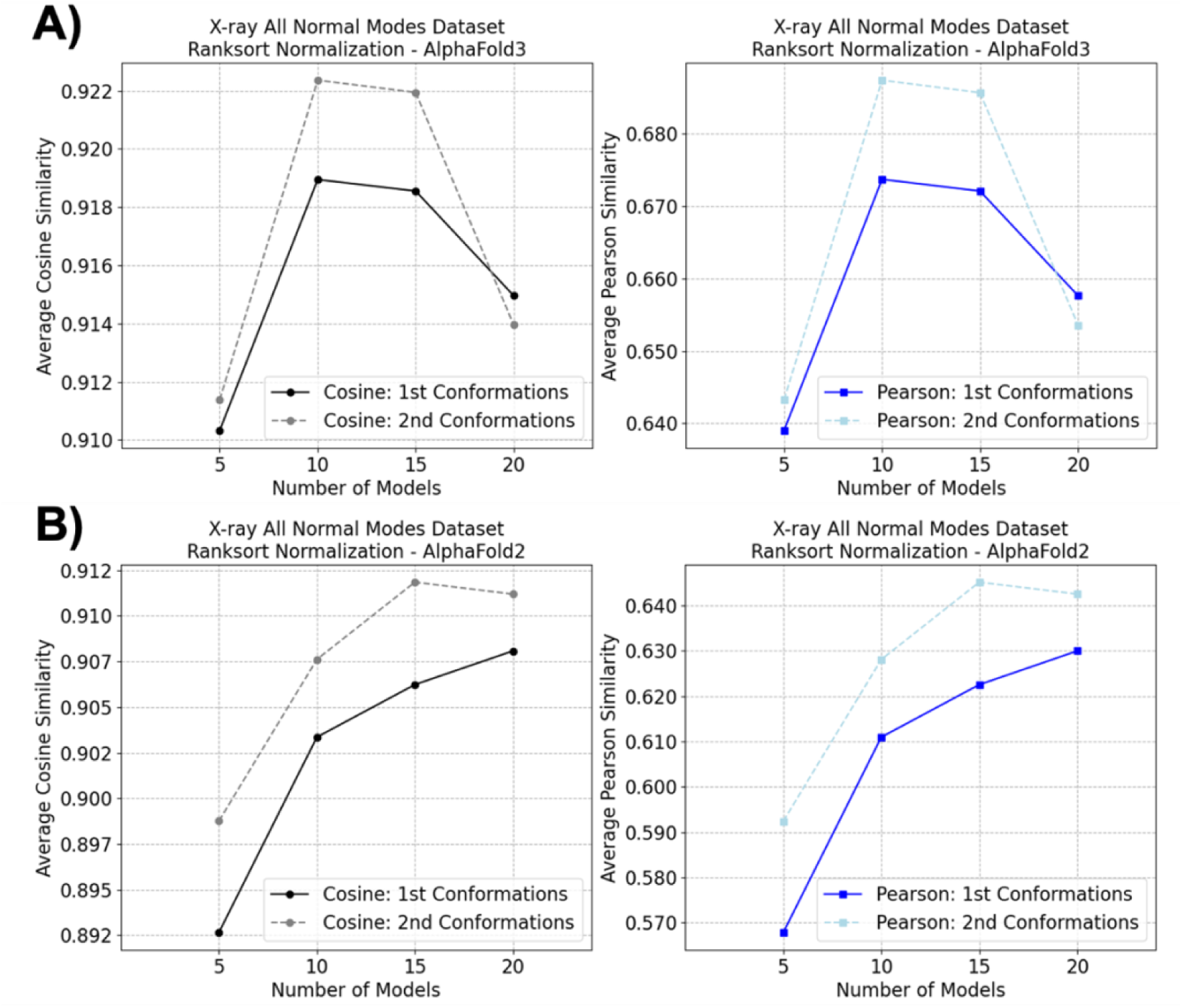
Impact of number of deep learning structural models on average similarity of X-ray all normal modes dataset, measured with cosine similarity (left panel, black and gray lines) and Pearson similarity (right panel, blue and light blue lines) for A) AlphaFold3 B) AlphaFold2. Continuous lines are for the first conformations set and dashed lines are for the second conformations set.

### MSF from Cryo-EM Dataset and Deep Learning Predicted Structural Models

Unfortunately, the cryo-EM dataset that we compiled do not contain multiple models of a protein. As a result, we are left only with the option of using normal mode ensembles as the reference MSF. As an example, all normal modes of 9yin protein were used for experimental MSF calculations and they were projected onto the protein structure after ranksort normalization (**Figure 7** A). Both projections and 2D plots of the MSF demonstrate a good agreement between the experimental and the deep learning ensemble MSF in terms of cosine similarity (**Figure 7** A and B) and Pearson similarity (**Figure S13** B). Average cosine similarity between the 82 experimental MSF and AlphaFold3 MSF is 0.926±0.031. It is 0.900±0.044 for AlphaFold2 and 0.873±0.045 for RosettaFold (**Figure 7** C and **Figure 8** A left panel). The average agreement is quite good for all deep learning structure prediction methods both in terms of cosine similarity (**Figure 8** A left panel) and Pearson similarity (**Figure 8** A right panel). Furthermore, we calculated MSF from the first 10 normal modes, which are the most collective modes (**Figure S14** and **Figure S15**). The normal mode MSF calculated with all modes and 10 lowest frequency modes produce similar results for AlphaFold3, AlphaFold2 and RosettaFold in terms of average cosine and Pearson similarities (**Figure 8** A and B). Taken together, all the cryo-EM results corroborate the central hypothesis of this work: deep learning predicted structural models can be used as proxies for protein flexibilities.

**Figure 7.**
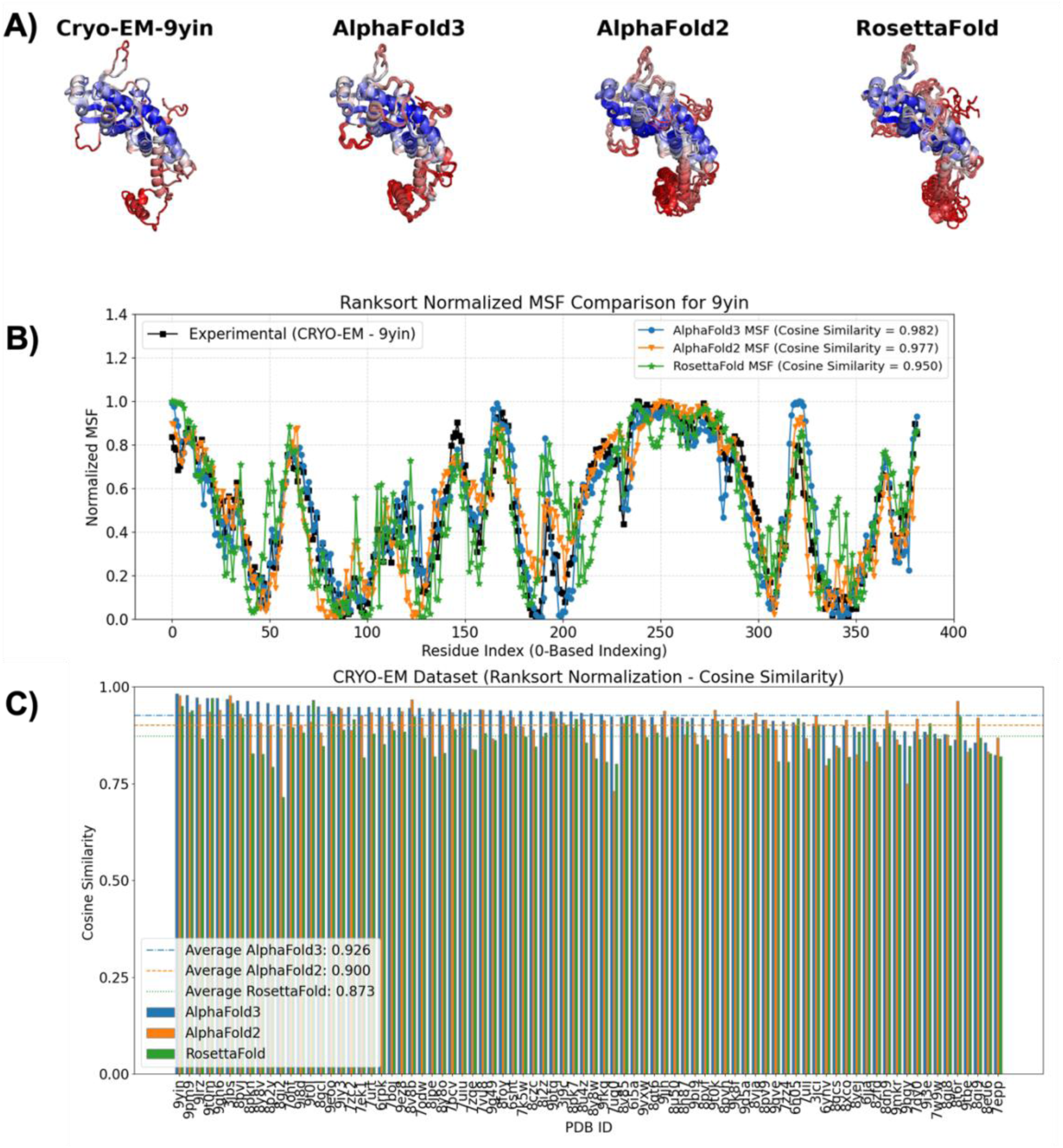
Ranksorted mean squared fluctuations (MSF) from all normal modes of cryo-EM structures and three deep learning structure prediction methods (AlphaFold3, AlphaFold2 and RosettaFold). A) Projections of the ranksorted MSF onto protein structures for 9yin. Blue-White-Red color palette is used for the projections, where blue indicates low flexibility and red indicates high flexibility. B) 2D comparison of the experimental and the computed MSF for 9yin. Black line (with squares) is for the experimental data, blue line (with circles) is for AlphaFold3, orange line (with inverse triangles) is for AlphaFold2 and green line (with stars) is for RosettaFold. C) Cosine similarity of the experimental and the computed MSF for 82 proteins in the cryo-EM dataset. AlphaFold3 bars are blue, AlphaFold2 bars are orange and RosettaFold bars are green. Averages of the cosine similarities over the entire dataset are also provided as horizontal lines for AlphaFold3 (blue dot-dashed line), AlphaFold2 (orange dashed line) and RosettaFold (green dotted line).

**Figure 8.**
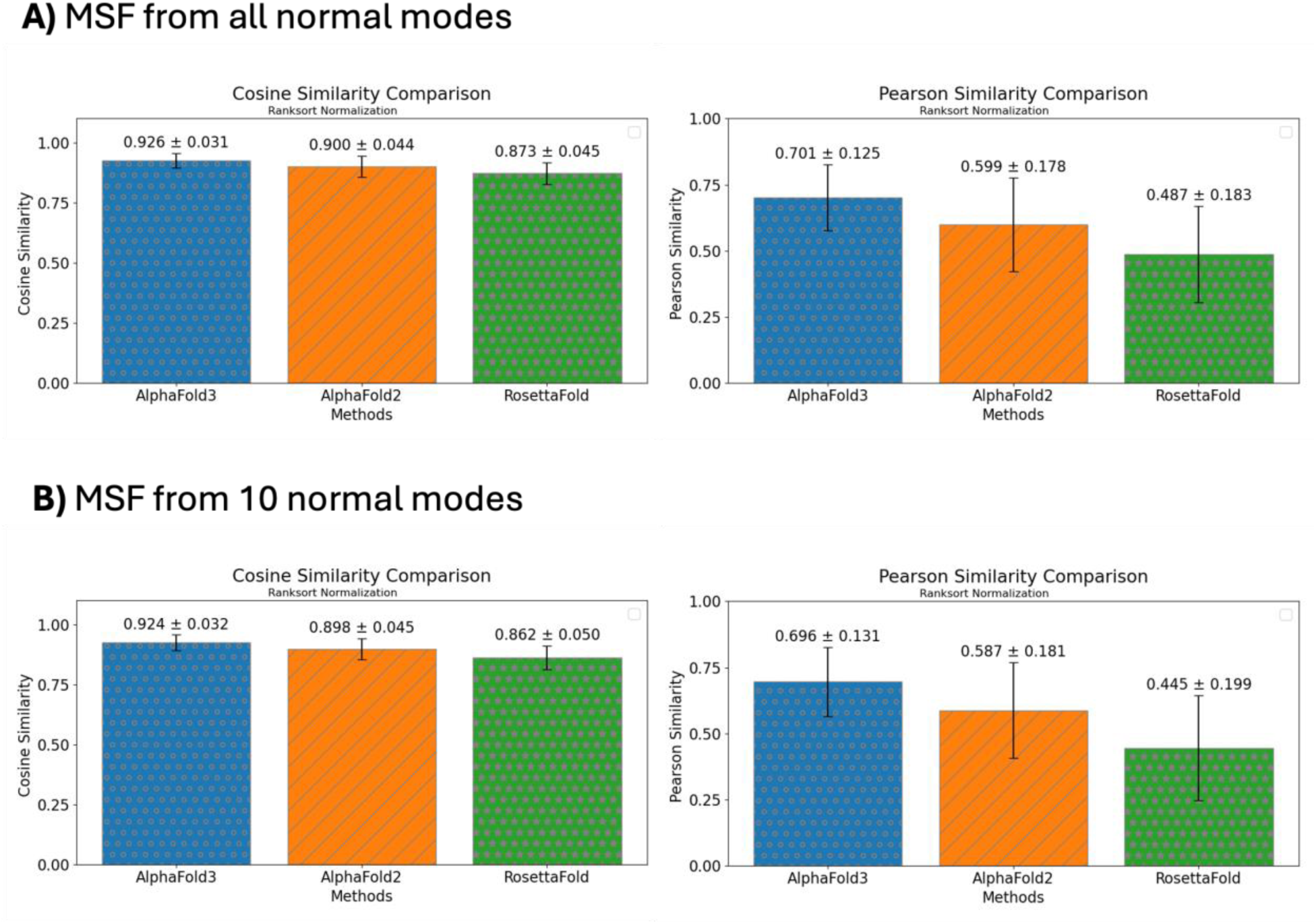
Average similarities with standard deviations as error bars for different approaches to obtain MSF from all normal modes of the cryo-EM experimental data. The average cosine similarities are given in the left panel and the average Pearson similarities are provided in the right panel. The average was taken over the similarities of 82 proteins and only Calpha atoms were used for normal mode analysis. Blue bars with gray circles are for AlphaFold3, orange bars with gray stripes are for AlphaFold2 and green bars with gray stars are for RosettaFold. A) All normal modes. B) 10 normal modes.

#### Impact of Number of Deep Learning Structural Models on the MSF of the Cryo-EM Dataset

Similar to the analyses for the NMR and X-ray datasets, we investigated effect of number models on the MSF prediction quality using cosine and Pearson similarities by using 5, 10, 15 and 20 deep learning structural models for the cryo-EM dataset. When all normal mode flexibilities are considered as the reference MSF, we observe an increase of the similarities when 15 or a smaller number of models from AlphaFold3 are employed (**Figure 9** A). The similarities start to deteriorate when 20 AlphaFold3 models are used. Even though the similarities still increase for 20 AlphaFold2 models, it is evident that the rate of increase starts to slow down (**Figure 9** B). The results for the lowest 10 normal modes are quite similar to the results based on all normal mode calculations (**Figure S16**).

**Figure 9.**
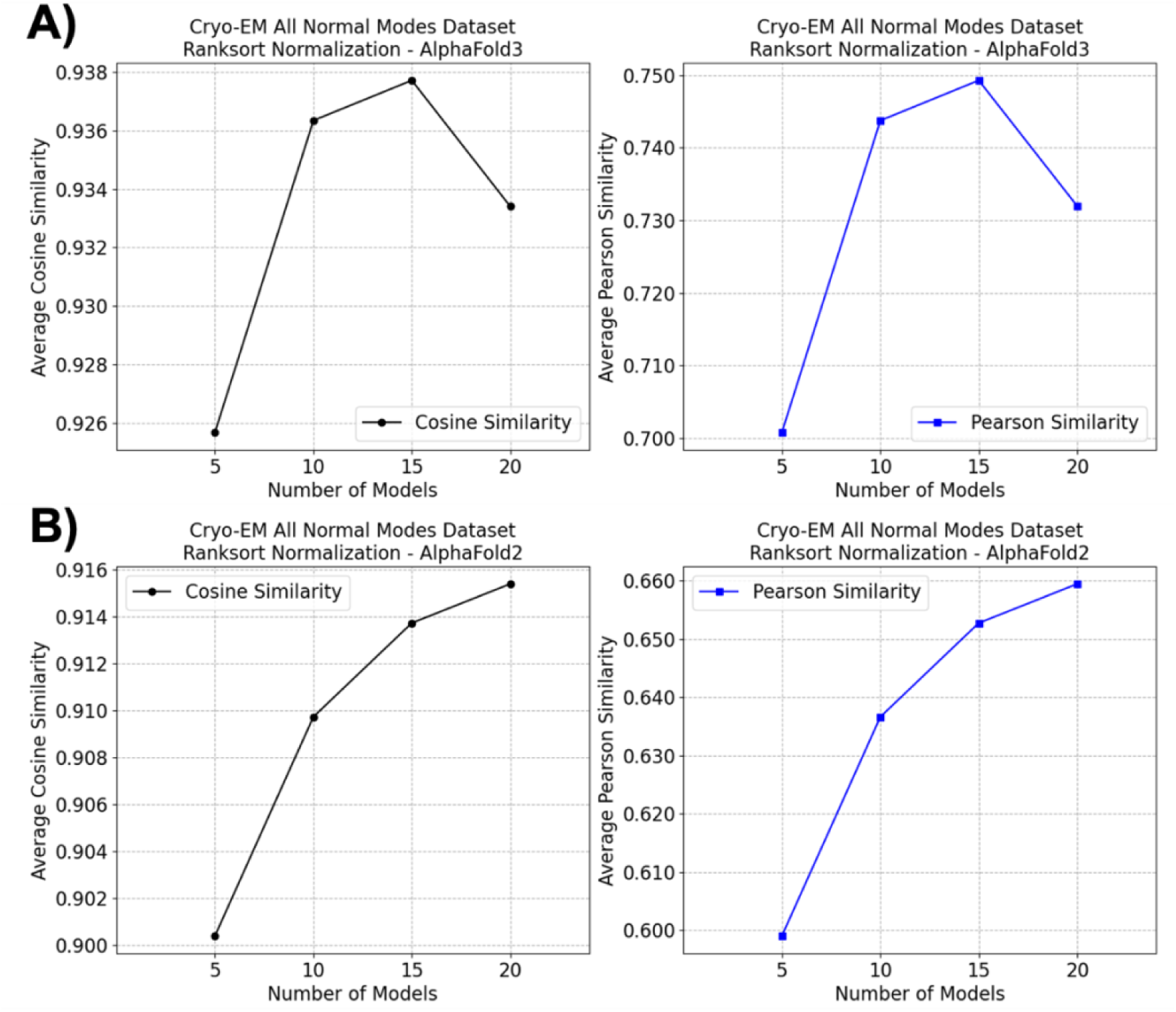
Impact of number of deep learning structural models on the cryo-EM all normal modes dataset, measured with cosine similarity (left panel) and Pearson similarity (right panel) for A) AlphaFold3 B) AlphaFold2.

### MSF from MD Simulations Dataset and Deep Learning Predicted Structural Models

MSF or its square root (RMSF) are one of the most widely used standard analyses in MD simulations of proteins. It quantifies protein flexibility from structural ensembles generated by the simulations. Here, we compared the MSF from 10 MD simulations with the MSF from deep learning structural ensembles of AlphaFold3 and AlphaFold2 methods. The MSF projections onto a reference structure for chain G of 6crk^57^ protein and 2D plot of it are presented in **Figure 10** A and B. The MSF from AlphaFold3 and AlphaFold2 demonstrate a varying degree of cosine similarity with different simulations. However, the lowest cosine similarity for AlphaFold3 is 0.904 (for simulation 2) and the lowest cosine similarity for AlphaFold2 is 0.889 (for simulation 2). Pearson similarities for this protein are presented in **Figure S17** A and B. Cosine similarities of each protein in the MD dataset are given **Figure 10** C, D and E. This analysis encounters a well-known limitation of the MD simulations: the sampling problem. The cosine correlation for simulation 1 of 3usy^58^ is quite high, while it reduces significantly in the second simulation. Despite this common problem, it is important to note that one can observe a decent agreement between the MSF from the deep learning structure prediction methods and the MSF from, at least, one of the MD simulations for all proteins in this dataset. The averages of the cosine and Pearson similarities over the entire dataset are also provided in **Figure 11**. AlphaFold3 is the best performer in MD simulations dataset as well. Overall, these results support the idea that the MSF from deep learning ensembles agree with the MSF obtained from established experimental data and/or simulated ensembles like MD simulations.

**Figure 10.**
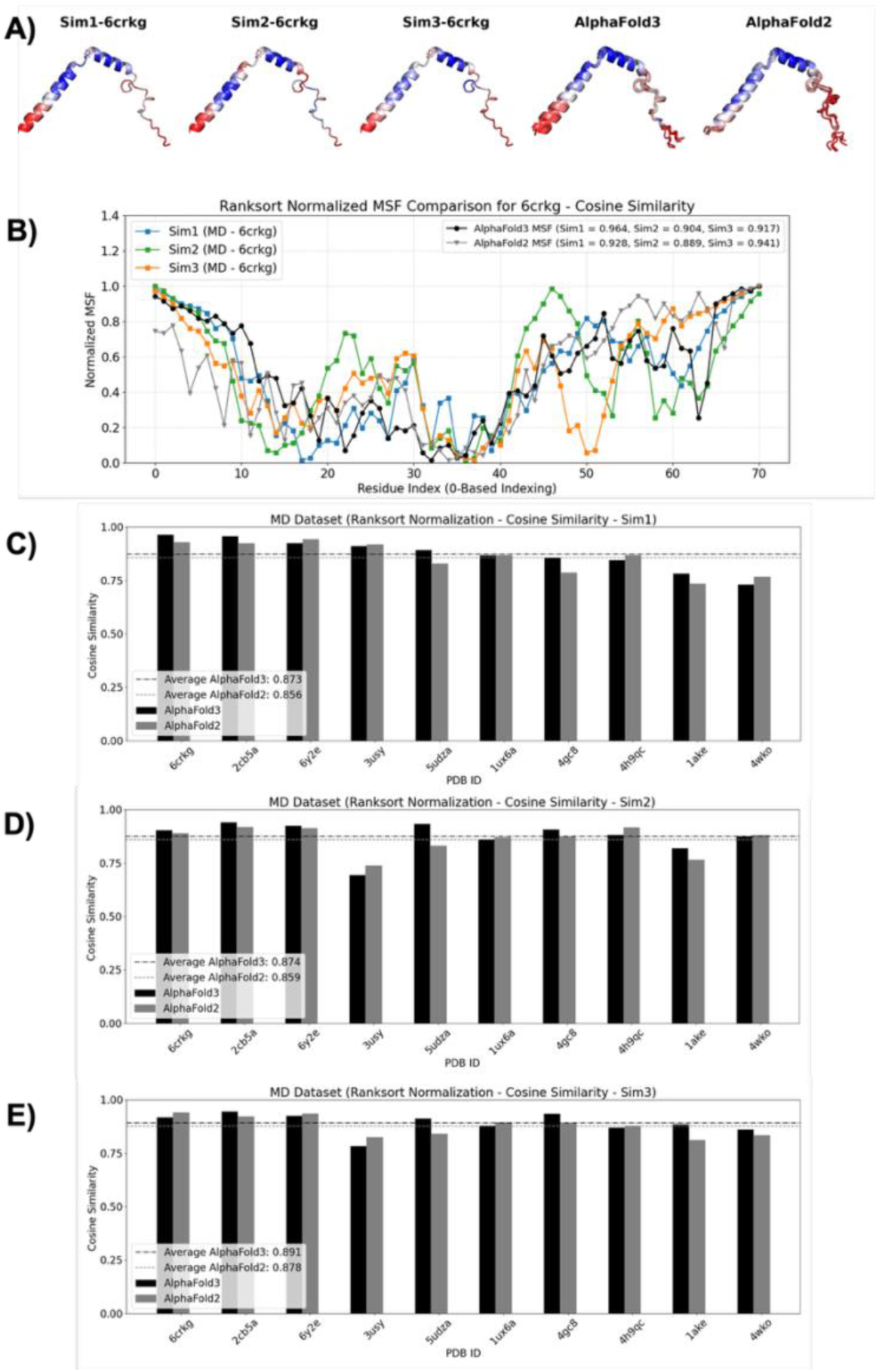
Ranksorted mean squared fluctuations (MSF) from molecular dynamics (MD) simulations and two deep learning structure prediction methods (AlphaFold3 and AlphaFold2). A) Projections of the ranksorted MSF onto protein structures for 6crk chain G. Blue-White-Red color palette is used for the projections, where blue indicates low flexibility and red indicates high flexibility. B) 2D comparison of the MD and the deep learning MSF for 6crk chain G. Blue (simulation 1), green (simulation 2) and orange (simulation 3) lines (with squares) are for the MD simulation data, black line (with circles) is for AlphaFold3, gray line (with inverse triangles) is for AlphaFold2. C) Cosine similarities of the first set of MD simulations and the deep learning ensemble MSF of 10 proteins in the MD dataset. D) Cosine similarities of the second set of MD simulations and the deep learning ensemble MSF of 10 proteins in the MD dataset. E) Cosine similarities of the third set of MD simulations and the deep learning ensemble MSF 10 proteins in the MD dataset. AlphaFold3 bars are black, AlphaFold2 bars are gray. Averages of the cosine similarities over the entire dataset are also provided as horizontal lines for AlphaFold3 (black dot-dashed line), AlphaFold2 (gray dashed line).

**Figure 11.**
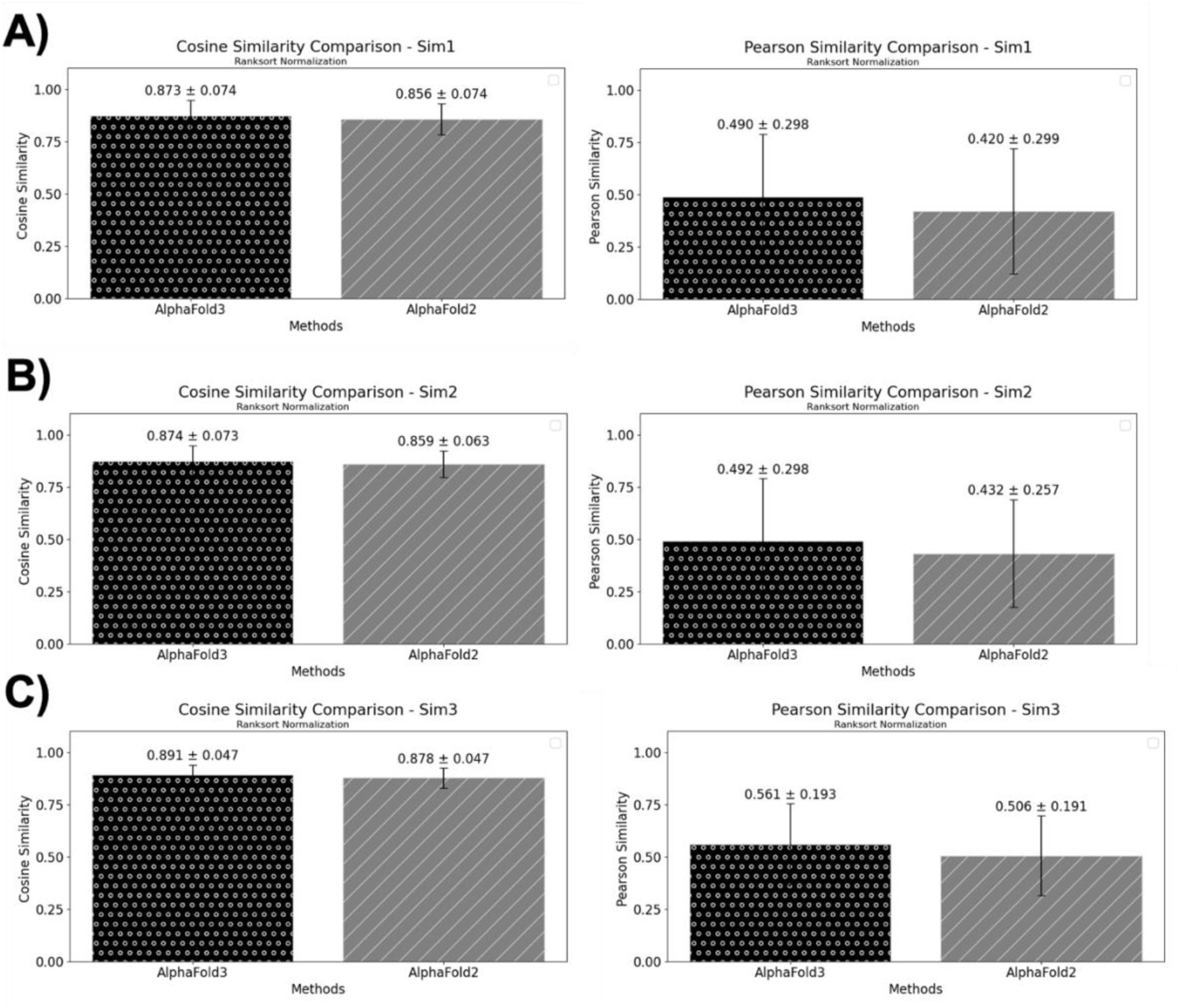
Average similarities with standard deviations as error bars for different approaches to obtain MSF from the molecular dynamics simulations dataset. The average cosine similarities are given in the left panel and the average Pearson similarities are provided in the right panel. The average was taken over the similarities of 10 proteins. Black bars with white circles are for AlphaFold3 and gray bars with white stripes are for AlphaFold2 A) Simulation set 1 (Sim1) B) Simulation set 2 (Sim2) C) Simulation set 3 (Sim3).

#### Impact of Number of Deep Learning Structural Models on the MSF of the MD Simulations Dataset

Similar to the previous analyses, we evaluated if increasing number of deep learning structural models would have an impact on the similarities between the deep learning ensemble MSF and a reference (MD simulations) ensemble MSF. We had observed that increasing number of models up to 15 improves the both the cosine similarity and Pearson similarity for the NMR, X-Ray and cryo-EM datasets. Interestingly, there is a decrease in the similarity metrics when more AlphaFold3 models are employed in the MSF calculations **Figure 12** A. It is more surprising to see that the situation is almost the opposite for the AlphaFold2 models (**Figure 12** B). AlphaFold2 trends are in agreement with our observations in the NMR, X-Ray and cryo-EM datasets.

**Figure 12.**
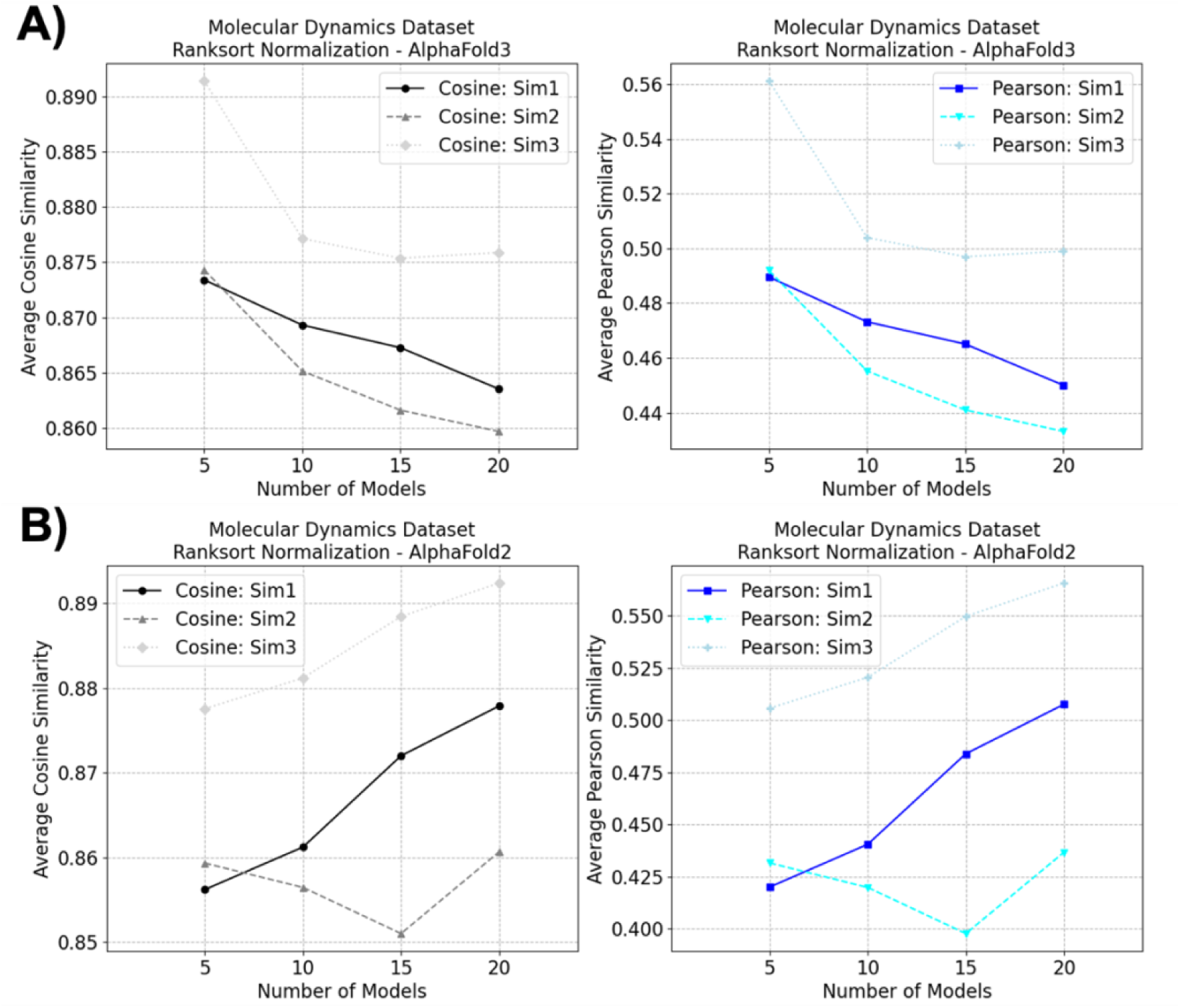
Impact of number of deep learning structural models on MSF similarities for the proteins in the molecular dynamics dataset, measured with cosine similarity (left panel) and Pearson similarity for A) AlphaFold3 B) AlphaFold2.

### Quantifying Terminal Effects on the Similarity of Deep Learning Ensemble MSF and Experimental/Simulated MSF

A previous study suggested that fluctuations in the terminal amino acids dominate the MSF calculations and the similarities performed based on them^59^. Even though we used rank-sorting to avoid this issue, we wondered if the fluctuations in N and C terminal residues are major contributors of the similarity we observed between the deep learning ensemble MSF and the experimental/simulated MSF. As a result, we systematically cropped MSF of 1, 2, 3, 4 and 5 amino acids from both terminals at the same time. Namely, when MSF of 5 amino acids cropped, we exclude 10 amino acids from the similarity computations. Then, we recalculated cosine and Pearson similarities for each protein in the NMR dataset (**Figure 13**). AlphaFold3 average cosine similarity slightly reduced from 0.940±0.034 to 0.927±0.040 while the average cosine similarity for AlphaFold2 changed from 0.915±0.041 to 0.899±0.046. RosettaFold cosine similarity values reduced from 0.918±0.040 to 0.901±0.048 (**Figure 13** A). Average Pearson similarities follow similar trends (**Figure 13** B). Even though there is a reduction, the reduction is not abrupt for any number of amino acid cropping and it is small. As a result, the data suggests that our results are robust even when terminal amino acids are not taken into account. The same analysis carried on X-Ray, cryo-EM and MD datasets support this conclusion (**Figures S18-S24**).

**Figure 13.**
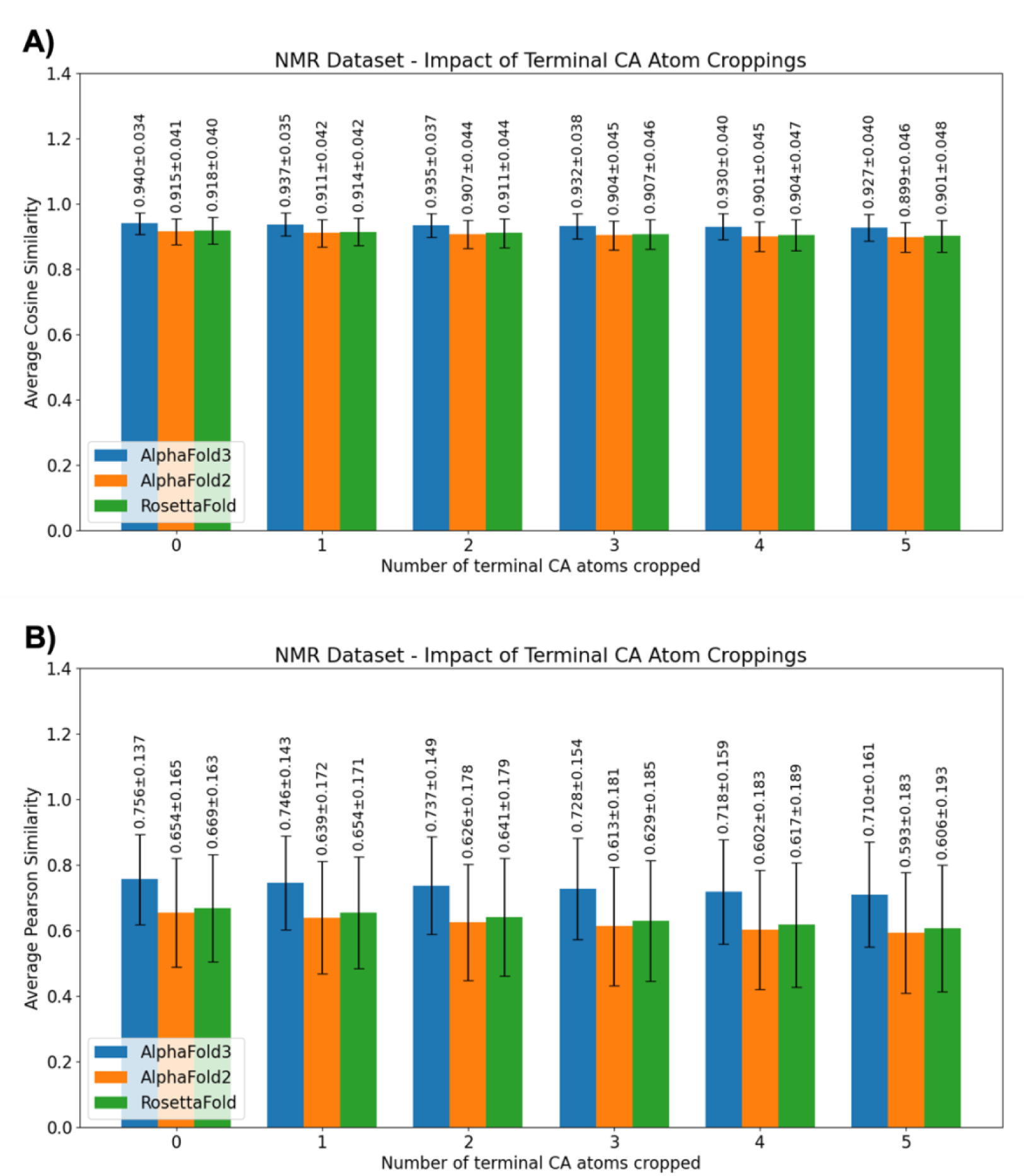
Systematic investigation of the impact of cropping MSF values of the terminal CA (Calpha) atoms for the NMR dataset. A) Average cosine similarity values B) Average Pearson similarity values.

### An Easy-to-use Cloud Interface for Calculating MSF from Deep Learning Predicted Structural Models

Up to this point, we tried to demonstrate that the MSF from the deep learning models agree with the experimental/simulated data. Here, we provide three easy-to-use Jupyter Notebooks on Google Cloud platform to calculate protein flexibilities for any protein from deep learning models, as described in this paper. The notebooks can be accessed at https://github.com/tekpinar/deepflex and they do not require any local installation. An interactive 2D plot of the MSF is generated with Plotly Python library^60^. Projections of the MSF onto the models are carried out with py3Dmol Python library ^61^. AlphaFold3 and AlphaFold2 MSF calculations require only zip files produced by the servers as input file, while RosettaFold notebook requests a multi-model single pdb file from the robetta server.

## CONCLUSIONS

In this work, we demonstrated that protein flexibilities, as measured by MSF, obtained from deep learning structural models agree, to a large extent, with three experimental datasets and a MD simulation dataset. We presented ranksorted MSF and compared similarity of the experimental and MD vs deep learning models with cosine and Pearson similarities. We observed that the flexibilities from AlphaFold3 models are generally better than AlphaFold2 and RosettaFold. We demonstrated robustness of our approach from different perspectives. Finally, we provided three publicly accessible notebooks for the scientific community to calculate MSF from deep learning-predicted structural models.

Despite the aforementioned problems of minmax normalization in flexibility analysis, we conducted minmax normalization to see if it can change the conclusions reached in this work. The results of minmax minimization are in respective pdf_files folders of NMR, X-Ray, cryo-EM and MD datasets in our zenodo dataset (https://doi.org/10.5281/zenodo.18823670). Importantly, our results demonstrate that the flexibility profiles derived from deep learning structural ensembles and experimental/simulated data remain in good agreement, regardless of the normalization procedure employed.

Furthermore, one should mention limitations of this study. All of the structures used in this study have less than 500 amino acids due to a limitation in the RosettaFold server (not the method itself). Even though we used two protein pairs (PDB IDs: 1d6m-1i7d and 1lfh-1lfg) with more than 500 amino acids, a large-scale study may be needed to reach a final conclusion about impact of protein size. We did not observe a significant correlation between protein size and the similarities for any dataset in this work. In addition to impact of protein size, we investigated if a correlation may be established between MSA depth and cosine/Pearson similarity metrics, using AlphaFold3 and AlphaFold2 generated MSAs. We were not able to observe any significant correlation.

Another limitation in this study is mainly monomeric state of the proteins. Effect of oligomerization on protein flexibilities has not been investigated. Unfortunately, majority of the NMR structures are small monomeric structures. Number of X-ray proteins of oligomeric structures in multiple conformational states is also limited. Due to these reasons, we focused only on monomeric cases and left the flexibilities of oligomeric structures to another investigation.

Our objective was to establish that deep learning structure prediction ensembles obtained from AlphaFold3, AlphaFold2 and RosettaFold can be used as proxies for protein flexibility, despite the fact that their initial aim was not to obtain protein flexibilities at all. Due to this reason, we excluded some recent methods such as BioEmu^62^, BioKinema^63^, SeaMoon^64^, OPUS-BFactor^65^ and PEGASUS^66^ targeting to obtain protein dynamics directly from sequence information.

Even though protein conformational sampling and protein flexibilities problems overlap partially, they should not be treated as identical problems. For example, normal modes of proteins are small fluctuations around an equilibrium, but they can determine protein flexibilities quite well. This point demonstrates that it is not necessary to obtain all conformational states of a protein to accurately determine its flexibility. Similarly, deep learning structural models do not necessarily have to contain all conformations of proteins to accurately describe protein flexibility, as demonstrated in this study.

To summarize, our results suggest that protein flexibilities, as measured by MSF, are substantially captured in deep learning structural ensembles, providing a computationally efficient proxy for flexibility analysis. AlphaFold3 is relatively the best algorithm in terms of average cosine and Pearson similarities to predict MSF. AlphaFold2 and RosettaFold follow it in terms of MSF prediction qualities. As the number of models increase up to 15 models, the flexibility prediction accuracy increases in general. Our results are consistent even if the terminal amino acids flexibilities are excluded in the similarity measurements. The results obtained in this work are important because they provide further evidence that protein dynamics may also be encoded in the sequence information. Finally, this study demonstrates that judicious utilization of existing deep learning algorithms can obviate the need for spending thousands of GPU/CPU-hours for training and testing new deep learning models or running compute-intensive methods for understanding protein flexibilities.

## DATA AND CODE AVAILABILITY

All data generated for this project is deposited at https://doi.org/10.5281/zenodo.18823670. The data includes input files, analyses, scripts and csv files for each experimental method. Furthermore, the repository includes 35 pdf files containing individual analyses of each protein from different perspectives. Furthermore, the authors provide three Google Colab notebooks at https://github.com/tekpinar/deepflex. The notebooks allow any user to calculate protein flexibilities from AlphaFold3, AlphaFold2 or RosettaFold outputs.

## ASSOCIATED CONTENT

## Supporting Information

The following file is available free of charge.

Supporting Information, which contains 24 figures and 4 tables (file type: PDF

**Figure S1.** Percentage identity matrix for A) 70 proteins in the NMR dataset (see Table S1 for further information) B) 43 proteins in the X-Ray dataset (see Table S2 for further information) C) 82 proteins in the cryo-EM dataset (see Table S3 for further information) D) 10 proteins in the molecular dynamics dataset (see Table S4 for further information).

**Figure S2.** Ranksorted mean squared fluctuations (MSF) from NMR method and three deep learning structure prediction methods (AlphaFold3, AlphaFold2 and RosettaFold). A) Projections of ranksorted MSF onto protein structures for 2lah. Blue-White-Red color palette is used for the projections, where blue indicates low flexibility and red indicates high flexibility. B) 2D comparison of the experimental and the computed MSF for 2ln3. Black line (with squares) is for experimental data, blue line (with circles) is for AlphaFold3, orange line (with inverse triangles) is for AlphaFold2 and green line (with stars) is for RosettaFold. C) Pearson similarity of experimental and computed MSF for 70 proteins in the NMR dataset. AlphaFold3 bars are blue, AlphaFold2 bars are orange and RosettaFold bars are green. Averages of the pearson similarities over the entire dataset are also provided as horizontal lines for AlphaFold3 (blue dot-dashed line), AlphaFold2 (orange dashed line) and RosettaFold (green dotted line).

**Figure S3.** Ranksorted mean squared fluctuations (MSF) from dual conformations of X-Ray structures and three deep learning structure prediction methods (AlphaFold3, AlphaFold2 and RosettaFold). A) Projections of the ranksorted MSF onto protein structures for 3fweB and 4hzfA. Blue-White-Red color palette is used for the projections, where blue indicates low flexibility and red indicates high flexibility. B) 2D comparison of the experimental and the computed MSF for 3fweB and 4hzfA. Black line (with squares) is for the experimental data, blue line (with circles) is for AlphaFold3, orange line (with inverse triangles) is for AlphaFold2 and green line (with stars) is for RosettaFold. C) Pearson similarity of the experimental and the computed MSF for 43 proteins in the X-Ray dataset. AlphaFold3 bars are blue, AlphaFold2 bars are orange and RosettaFold bars are green. Averages of the Pearson similarities over the entire dataset are also provided as horizontal lines for AlphaFold3 (blue dot-dashed line), AlphaFold2 (orange dashed line) and RosettaFold (green dotted line).

**Figure S4.** Ranksorted mean squared fluctuations (MSF) from dual conformations of X-Ray structures and three deep learning structure prediction methods (AlphaFold3, AlphaFold2 and RosettaFold) for two proteins with more than 500 amino acids. A) Projections of the ranksorted MSF onto protein structures for 1d6m-1i7d. Blue-White-Red color palette is used for the projections, where blue indicates low flexibility and red indicates high flexibility. B) 2D comparison of the experimental and the computed MSF for 1d6m-1i7d protein pair. Black line (with squares) is for the experimental data, blue line (with circles) is for AlphaFold3 and orange line (with inverse triangles) is for AlphaFold2. C) Projections of the ranksorted MSF onto protein structures for 1lfh-1lfg. D) 2D comparison of the experimental and the computed MSF for 1lfh-1lfg protein pair. Black line (with squares) is for the experimental data, blue line (with circles) is for AlphaFold3 and orange line (with inverse triangles) is for AlphaFold2.

**Figure S5.** Ranksorted mean squared fluctuations (MSF) from <u>all normal modes of X-Ray structures</u> and three deep learning structure prediction methods (AlphaFold3, AlphaFold2 and RosettaFold). A) Projections of the ranksorted MSF onto protein structures for 1nfk chain A and 1ooa chain A. Blue-White-Red color palette is used for the projections, where blue indicates low flexibility and red indicates high flexibility. B) 2D comparison of the experimental and the computed MSF for 1nfk chain A and 1ooa chain A. Black line (with squares) is for the experimental data, blue line (with circles) is for AlphaFold3, orange line (with inverse triangles) is for AlphaFold2 and green line (with stars) is for RosettaFold. C) Cosine similarities of the experimental and the computed MSF for the first conformation of 43 proteins in the X-Ray dataset. AlphaFold3 bars are blue, AlphaFold2 bars are orange and RosettaFold bars are green. Averages of the cosine similarities over the entire dataset are also provided as horizontal lines for AlphaFold3 (blue dot-dashed line), AlphaFold2 (orange dashed line) and RosettaFold (green dotted line). D) Cosine similarities of the experimental and the computed MSF for the second conformation of 43 proteins in the X-Ray dataset. The colors are the same as in C.

**Figure S6.** Ranksorted mean squared fluctuations (MSF) from <u>all normal modes of X-Ray structures</u> and three deep learning structure prediction methods (AlphaFold3, AlphaFold2 and RosettaFold). A) Projections of the ranksorted MSF onto protein structures for 1nfk chain A and 1ooa chain A. Blue-White-Red color palette is used for the projections, where blue indicates low flexibility and red indicates high flexibility. B) 2D comparison of the experimental and the computed MSF for 1nfk chain A and 1ooa chain A. Black line (with squares) is for the experimental data, blue line (with circles) is for AlphaFold3, orange line (with inverse triangles) is for AlphaFold2 and green line (with stars) is for RosettaFold. C) Pearson similarities of the experimental and the computed MSF for the first conformation of 43 proteins in the X-Ray dataset. AlphaFold3 bars are blue, AlphaFold2 bars are orange and RosettaFold bars are green. Averages of Pearson similarities over the entire dataset are also provided as horizontal lines for AlphaFold3 (blue dot-dashed line), AlphaFold2 (orange dashed line) and RosettaFold (green dotted line). D) Pearson similarities of the experimental and the computed MSF for the second conformation of 43 proteins in the X-Ray dataset. The colors are the same as in C.

**Figure S7.** Ranksorted mean squared fluctuations (MSF) from <u>the lowest frequency 10 normal modes of X-Ray structures</u> and three deep learning structure prediction methods (AlphaFold3, AlphaFold2 and RosettaFold). A) Projections of the ranksorted MSF onto protein structures for 2jej chain A and 3fds chain A. Blue-White-Red color palette is used for the projections, where blue indicates low flexibility and red indicates high flexibility. B) 2D comparison of the experimental and the computed MSF for 2jej chain A and 3fds chain A. Black line (with squares) is for the experimental data, blue line (with circles) is for AlphaFold3, orange line (with inverse triangles) is for AlphaFold2 and green line (with stars) is for RosettaFold. C) Cosine similarities of the experimental and the computed MSF for the first conformation of 43 proteins in the X-Ray dataset. AlphaFold3 bars are blue, AlphaFold2 bars are orange and RosettaFold bars are green. Averages of the cosine similarities over the entire dataset are also provided as horizontal lines for AlphaFold3 (blue dot-dashed line), AlphaFold2 (orange dashed line) and RosettaFold (green dotted line). D) Cosine similarities of the experimental and the computed MSF for the second conformation of 43 proteins in the X-Ray dataset. The colors are the same as in C.

**Figure S8.** Ranksorted mean squared fluctuations (MSF) from <u>the lowest frequency 10 normal modes of X-Ray structures</u> and three deep learning structure prediction methods (AlphaFold3, AlphaFold2 and RosettaFold). A) Projections of the ranksorted MSF onto protein structures for 2jej chain A and 3fds chain A. Blue-White-Red color palette is used for the projections, where blue indicates low flexibility and red indicates high flexibility. B) 2D comparison of the experimental and the computed MSF for 2jej chain A and 3fds chain A. Black line (with squares) is for the experimental data, blue line (with circles) is for AlphaFold3, orange line (with inverse triangles) is for AlphaFold2 and green line (with stars) is for RosettaFold. C) Pearson similarities of the experimental and the computed MSF for the first conformation of 43 proteins in the X-Ray dataset. AlphaFold3 bars are blue, AlphaFold2 bars are orange and RosettaFold bars are green. Averages of Pearson similarities over the entire dataset are also provided as horizontal lines for AlphaFold3 (blue dot-dashed line), AlphaFold2 (orange dashed line) and RosettaFold (green dotted line). D) Pearson similarities of the experimental and the computed MSF for the second conformation of 43 proteins in the X-Ray dataset. The colors are the same as in C.

**Figure S9.** Averages of the similarities for different approaches to obtain MSF from the second conformation set of the X-Ray experimental data. The average cosine similarities are given in the left panel and the average Pearson similarities are provided in the right panel. The average was taken over the similarities of 43 proteins. Blue bars with gray circles are for AlphaFold3, orange bars with gray stripes are for AlphaFold2 and green bars with gray stars are for RosettaFold. A) Two experimental protein conformations were used for the MSF calculations. B) All normal modes of the second conformations was used for the MSF calculations. Only Calpha atoms were used for normal mode analysis. C) Only 10 lowest eigenvalue normal modes of each conformation were used for normal mode analysis. Only Calpha atoms were used for normal mode analysis. D) Bfactors were ranksort normalized and they were used as a proxy for the MSF.

**Figure S10.** Impact of number of deep learning structural models on average similarity of X-ray 10 normal modes dataset, measured with cosine similarity (left panel, black and gray lines) and Pearson similarity (right panel, blue and light blue lines) for A) AlphaFold3 B) AlphaFold2. Continuous lines are for the first conformations set and dashed lines are for the second conformations set.

**Figure S11.** Impact of number of deep learning structural models on average similarity of X-ray dual conformations dataset, measured with cosine similarity (left panel, black line) and Pearson similarity (right panel, blue line) for A) AlphaFold3 B) AlphaFold2.

**Figure S12.** Impact of number of deep learning structural models on average similarity of X-ray bfactors dataset, measured with cosine similarity (left panel, black and gray lines) and Pearson similarity (right panel, blue and light blue lines) for A) AlphaFold3 B) AlphaFold2. Continuous lines are for the first conformations set and dashed lines are for the second conformations set.

**Figure S13.** Ranksorted mean squared fluctuations (MSF) from <u>all normal modes</u> of the cryo-EM structures and three deep learning structure prediction methods (AlphaFold3, AlphaFold2 and RosettaFold). A) Projections of the ranksorted MSF onto protein structures for 9yin. Blue-White-Red color palette is used for the projections, where blue indicates low flexibility and red indicates high flexibility. B) 2D comparison of the experimental and the computed MSF for 9yin. Black line (with squares) is for the experimental data, blue line (with circles) is for AlphaFold3, orange line (with inverse triangles) is for AlphaFold2 and green line (with stars) is for RosettaFold. C) Pearson similarity of the experimental and the computed MSF for 82 proteins in the cryo-EM dataset. AlphaFold3 bars are blue, AlphaFold2 bars are orange and RosettaFold bars are green. Averages of the Pearson similarities over the entire dataset are also provided as horizontal lines for AlphaFold3 (blue dot-dashed line), AlphaFold2 (orange dashed line) and RosettaFold (green dotted line)

**Figure S14.** Ranksorted mean squared fluctuations (MSF) from <u>the first 10 normal modes</u> of the cryo-EM structures and three deep learning structure prediction methods (AlphaFold3, AlphaFold2 and RosettaFold). A) Projections of the ranksorted MSF onto protein structures for 9pm9. Blue-White-Red color palette is used for the projections, where blue indicates low flexibility and red indicates high flexibility. B) 2D comparison of the experimental and the computed MSF for 9pm9. Black line (with squares) is for the experimental data, blue line (with circles) is for AlphaFold3, orange line (with inverse triangles) is for AlphaFold2 and green line (with stars) is for RosettaFold. C) Cosine similarity of the experimental and the computed MSF for 82 proteins in the cryo-EM dataset. AlphaFold3 bars are blue, AlphaFold2 bars are orange and RosettaFold bars are green. Averages of the cosine similarities over the entire dataset are also provided as horizontal lines for AlphaFold3 (blue dot-dashed line), AlphaFold2 (orange dashed line) and RosettaFold (green dotted line).

**Figure S15.** Ranksorted mean squared fluctuations (MSF) from <u>the first 10 normal modes</u> of the cryo-EM structures and three deep learning structure prediction methods (AlphaFold3, AlphaFold2 and RosettaFold). A) Projections of the ranksorted MSF onto protein structures for 9pm9. Blue-White-Red color palette is used for the projections, where blue indicates low flexibility and red indicates high flexibility. B) 2D comparison of the experimental and the computed MSF for 9pm9. Black line (with squares) is for the experimental data, blue line (with circles) is for AlphaFold3, orange line (with inverse triangles) is for AlphaFold2 and green line (with stars) is for RosettaFold. C) Pearson similarity of the experimental and the computed MSF for 82 proteins in the cryo-EM dataset. AlphaFold3 bars are blue, AlphaFold2 bars are orange and RosettaFold bars are green. Averages of the Pearson similarities over the entire dataset are also provided as horizontal lines for AlphaFold3 (blue dot-dashed line), AlphaFold2 (orange dashed line) and RosettaFold (green dotted line).

**Figure S16.** Impact of number of deep learning structural models on the cryo-EM 10 normal modes dataset, measured with cosine similarity (left panel) and Pearson similarity (right panel) for A) AlphaFold3 B) AlphaFold2.

**Figure S17.** Ranksorted mean squared fluctuations (MSF) from molecular dynamics (MD) simulations and three deep learning structure prediction methods (AlphaFold3 and AlphaFold2). A) Projections of the ranksorted MSF onto protein structures for 6crk chain G. Blue-White-Red color palette is used for the projections, where blue indicates low flexibility and red indicates high flexibility. B) 2D comparison of the MD and the deep learning ensemble MSF for 6crk chain G. Blue (simulation 1), green (simulation 2) and orange (simulation 3) lines (with squares) are for the MD simulation data, black line (with circles) is for AlphaFold3, gray line (with inverse triangles) is for AlphaFold2. C) Pearson similarities of the first set of MD simulations and the deep learning ensemble MSF of 10 proteins in the MD dataset. D) Pearson similarities of the second set of MD simulations and the deep learning ensemble MSF of 10 proteins in the MD dataset. E) Pearson similarities of the third set of MD simulations and the deep learning ensemble MSF 10 proteins in the MD dataset. AlphaFold3 bars are black, AlphaFold2 bars are gray. Averages of the cosine similarities over the entire dataset are also provided as horizontal lines for AlphaFold3 (black dot-dashed line), AlphaFold2 (gray dashed line).

**Figure S18.** Systematic investigation of the impact of cropping MSF values of the terminal CA (Calpha) atoms for the X-Ray dual conformations dataset. A) Average cosine similarity values B) Average Pearson similarity values.

**Figure S19.** Systematic investigation of the impact of cropping MSF values of the terminal CA (Calpha) atoms for the X-Ray all normal modes dataset. The left panel is for the first conformations and the right panel is for the second conformations. A) Average cosine similarity values B) Average Pearson similarity values.

**Figure S20.** Systematic investigation of the impact of cropping MSF values of the terminal CA (Calpha) atoms for the X-Ray 10 normal modes dataset. The left panel is for the first conformations and the right panel is for the second conformations. A) Average cosine similarity values B) Average Pearson similarity values.

**Figure S21.** Systematic investigation of the impact of cropping MSF values of the terminal CA (Calpha) atoms for the X-Ray bfactors dataset. The left panel is for the first conformations and the right panel is for the second conformations. A) Average cosine similarity values B) Average Pearson similarity values.

**Figure S22.** Systematic investigation of the impact of cropping MSF values of the terminal CA (Calpha) atoms for the cryo-EM all normal modes dataset. A) Average cosine similarity values B) Average Pearson similarity values.

**Figure S23.** Systematic investigation of the impact of cropping MSF values of the terminal CA (Calpha) atoms for the cryo-EM 10 normal modes dataset. A) Average cosine similarity values B) Average Pearson similarity values

**Figure S24.** Systematic investigation of the impact of cropping MSF values of the terminal CA (Calpha) atoms for the molecular dynamics (MD) dataset. The left panel is for the first simulation set, the middle panel is for the second simulation set and the right panel is for the third simulation set. A) Average cosine similarity values B) Average Pearson similarity values.

**Table S1.** Protein Databank IDs, number of models and number of residues for all proteins in the NMR dataset. The proteins are ordered according to the number of residues. Three proteins highlighted with red text color have 10 models in their structures. All the remaining proteins have 20 models.

**Table S2.** Protein Databank IDs of two conformations (as PDB ID1 and PDB ID2), number of residues and RMSD between the conformations for all proteins in the X-Ray dataset. The fifth uppercase character in each PDB ID denotes the relevant chain. Please note that number of residues may have been modified to precisely match the pairs for some cases.

**Table S3.** Protein Databank IDs and number of residues for all proteins in the cryo-EM dataset.

**Table S4.** Protein Databank IDs, number of residues, number of frames and simulation times for all proteins in the molecular dynamics simulations dataset. The fifth uppercase character in each PDB ID denotes the relevant chain.

## AUTHOR INFORMATION

### Author Contributions

The manuscript was written through contributions of all authors. All authors have given approval to the final version of the manuscript.

### Notes

The authors declare no competing interests.

## ACKNOWLEDGMENT

The authors thank Google Deepmind for providing AlphaFold3 weights. The numerical calculations reported in this study were partially performed using the EuroHPC Joint Undertaking (EuroHPC JU) supercomputer MareNostrum 5, hosted by the Barcelona Supercomputing Center (BSC). The computational time was provided for the project entitled “Computing Structure and Multiple Sequence Alignments of All Human Protein Isoforms with Colabfold for Proteome Scale Analyses of Human Mutations (HumanIsoVar)”. Access to MareNostrum 5 was provided through a national access call coordinated by the Scientific and Technological Research Council of Turkey (TÜBİTAK). We gratefully acknowledge BSC, TÜBİTAK, and the EuroHPC JU for providing access to these resources and supporting this research. The authors used Gemini AI (v3) tool to generate <u>only and only</u> abstract of the text. The abstract was read and verified by the authors for scientific accuracy. The graphical abstract was generated by ChatGPT. The authors thank Prof. Ahmet YILDIRIM for critical reading of the manuscript.

## ABBREVIATIONS

ANM: Anisotropic Network Model
Cryo-EM: Cryo-Electron Microscopy
MD: Molecular Dynamics
MSA: Multiple Sequence Alignment
MSF: Mean Squared Fluctuations
NMA: Normal Mode Analysis
NMR: Nuclear Magnetic Resonance
PDB: Protein Databank
pLDDT: predicted Local Distance Difference Test
RMSD: Root Mean Squared Distance.

