## Supporting Information for "Deep Learning Structural Ensembles as Proxies for Protein Flexibility"

**This Supporting Information contains 24 figures and 4 tables.**

SUPPORTING FIGURES.

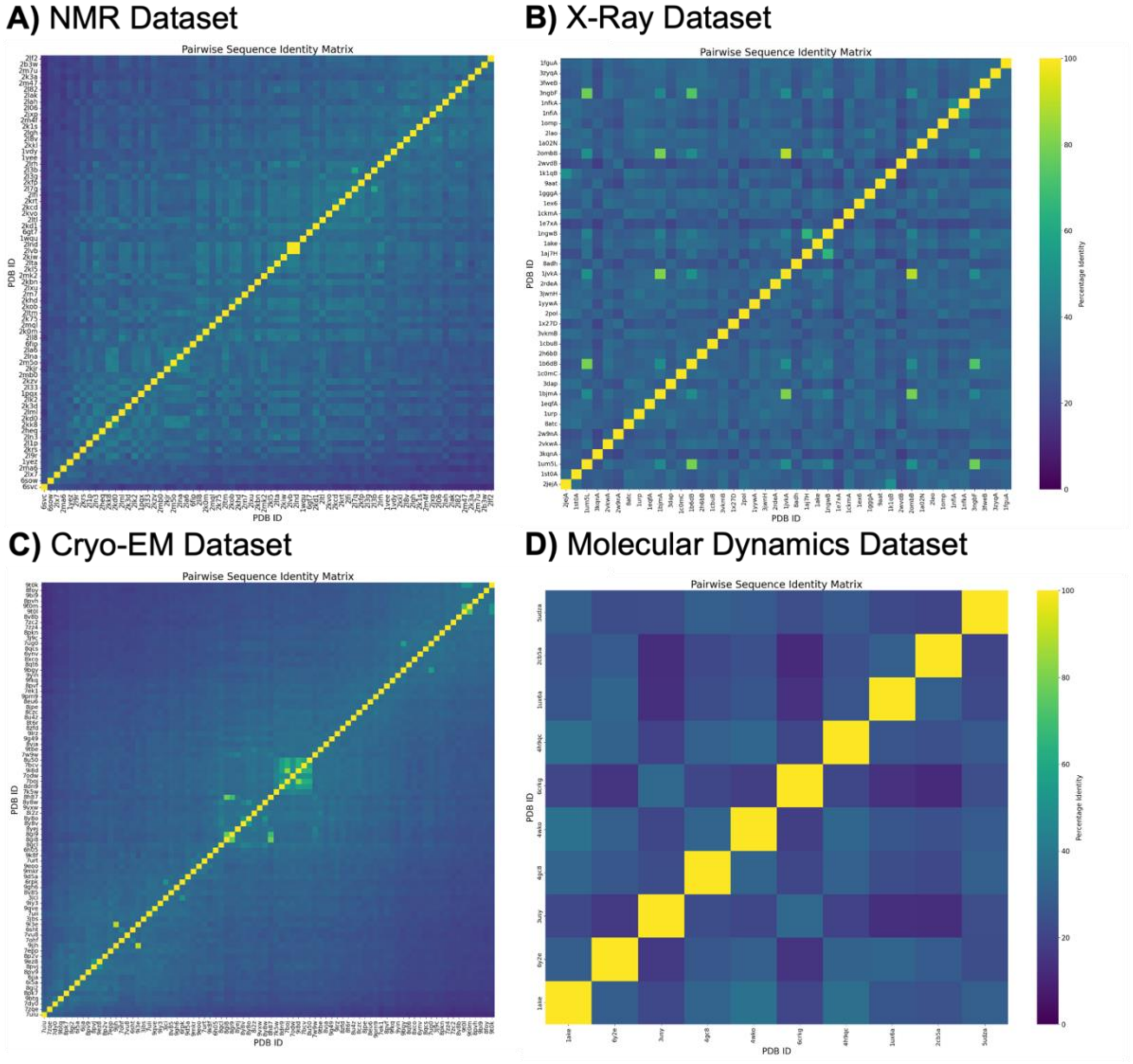


**Figure S1.** Percentage identity matrix for A) 70 proteins in the NMR dataset (see Table S1 for further information) B) 43 proteins in the X-Ray dataset (see Table S2 for further information) C) 82 proteins in the cryo-EM dataset (see Table S3 for further information) D) 10 proteins in the molecular dynamics dataset (see Table S4 for further information).

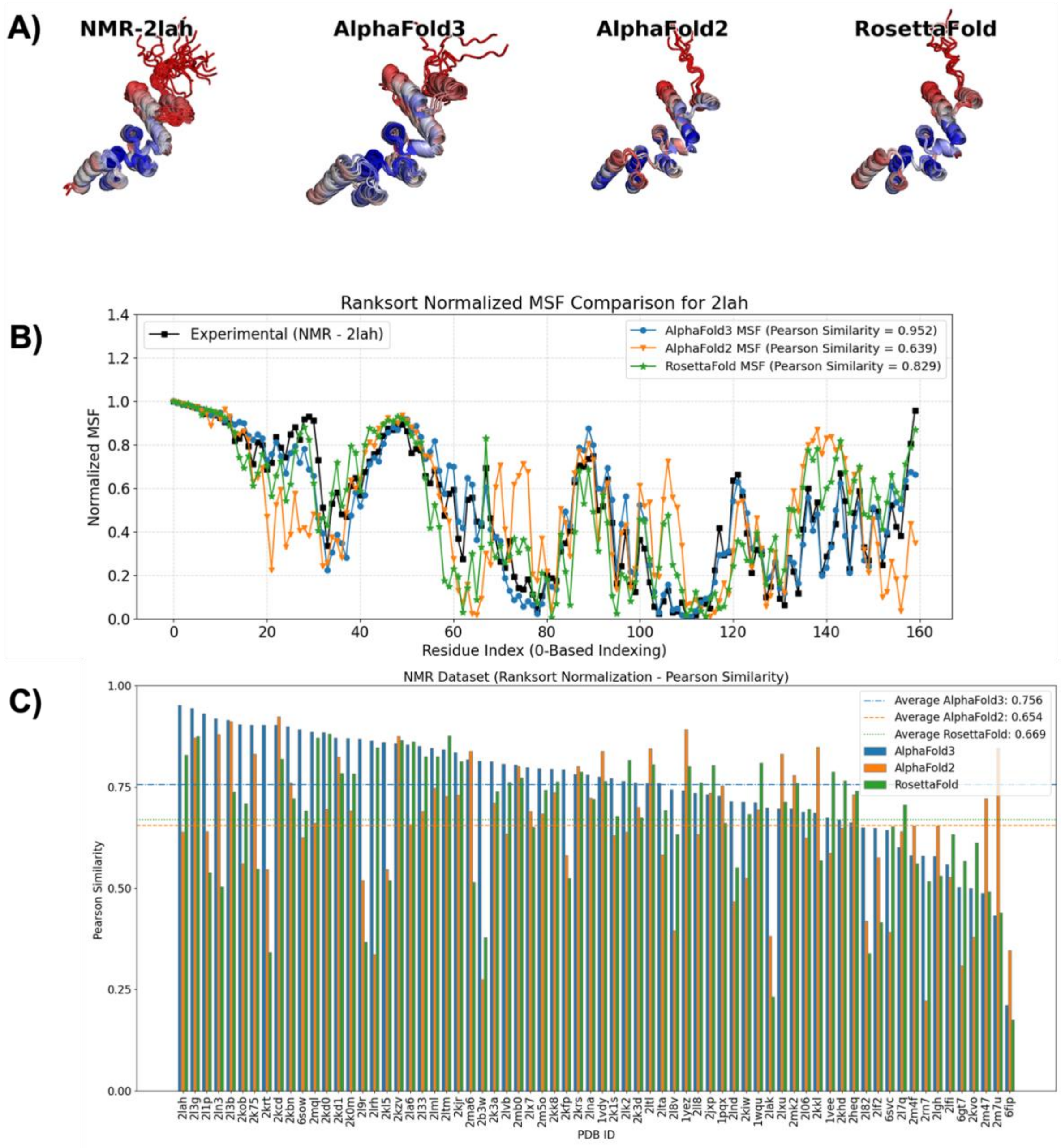


**Figure S2.** Ranksorted mean squared fluctuations (MSF) from NMR method and three deep learning structure prediction methods (AlphaFold3, AlphaFold2 and RosettaFold). A) Projections of ranksorted MSF onto protein structures for 2lah. Blue-White-Red color palette is used for the projections, where blue indicates low flexibility and red indicates high flexibility. B) 2D comparison of the experimental and the computed MSF for 2ln3. Black line (with squares) is for experimental data, blue line (with circles) is for AlphaFold3, orange line (with inverse triangles) is for AlphaFold2 and green line (with stars) is for RosettaFold. C) Pearson similarity of experimental and computed MSF for 70 proteins in the NMR dataset. AlphaFold3 bars are blue, AlphaFold2 bars are orange and RosettaFold bars are green. Averages of the pearson similarities over the entire dataset are also provided as horizontal lines for AlphaFold3 (blue dot-dashed line), AlphaFold2 (orange dashed line) and RosettaFold (green dotted line).

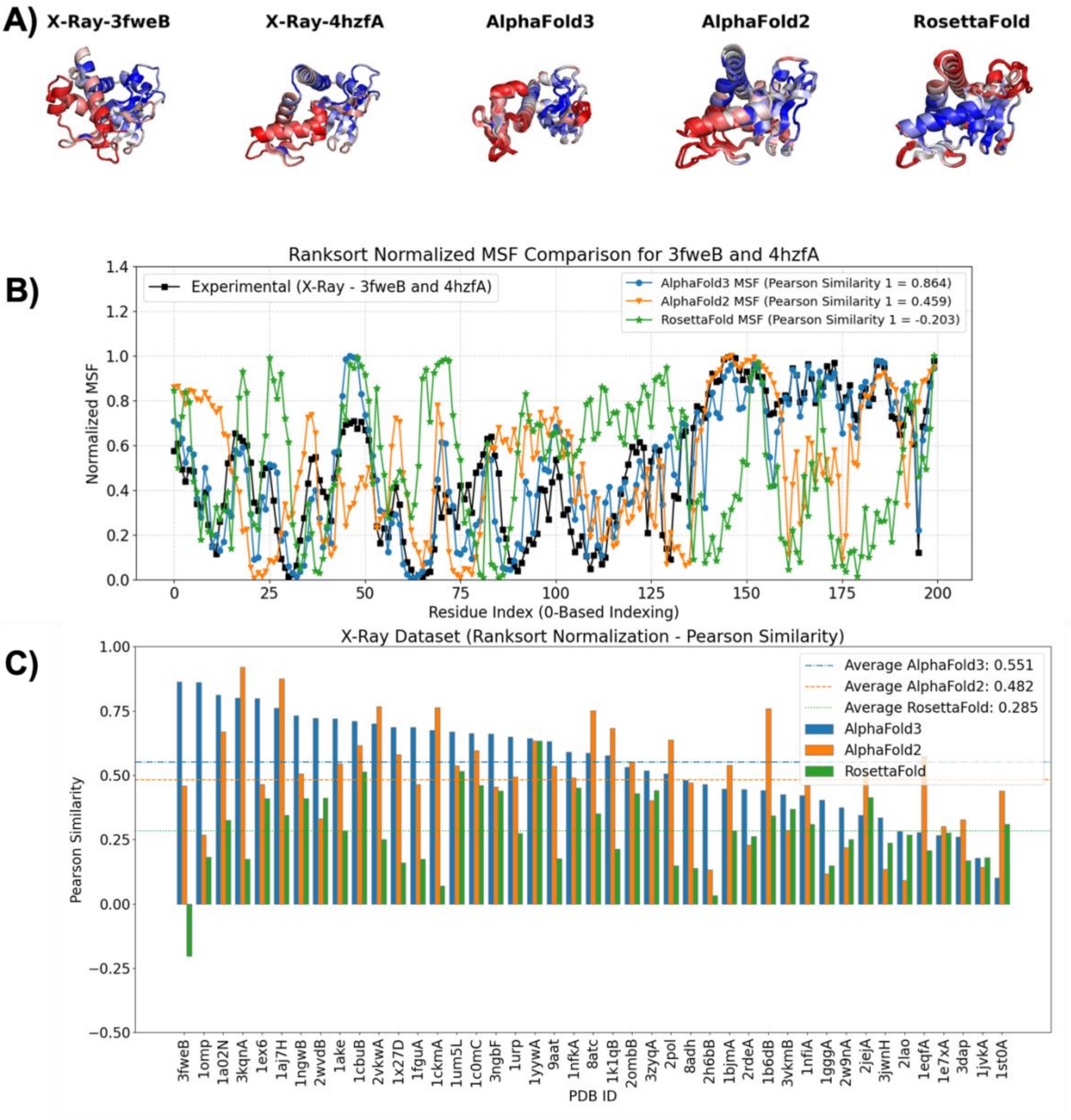


**Figure S3.** Ranksorted mean squared fluctuations (MSF) from dual conformations of X-Ray structures and three deep learning structure prediction methods (AlphaFold3, AlphaFold2 and RosettaFold). A) Projections of the ranksorted MSF onto protein structures for 3fweB and 4hzfA. Blue-White-Red color palette is used for the projections, where blue indicates low flexibility and red indicates high flexibility. B) 2D comparison of the experimental and the computed MSF for 3fweB and 4hzfA. Black line (with squares) is for the experimental data, blue line (with circles) is for AlphaFold3, orange line (with inverse triangles) is for AlphaFold2 and green line (with stars) is for RosettaFold. C) Pearson similarity of the experimental and the computed MSF for 43 proteins in the X-Ray dataset. AlphaFold3 bars are blue, AlphaFold2 bars are orange and RosettaFold bars are green. Averages of the Pearson similarities over the entire dataset are also provided as horizontal lines for AlphaFold3 (blue dot-dashed line), AlphaFold2 (orange dashed line) and RosettaFold (green dotted line).

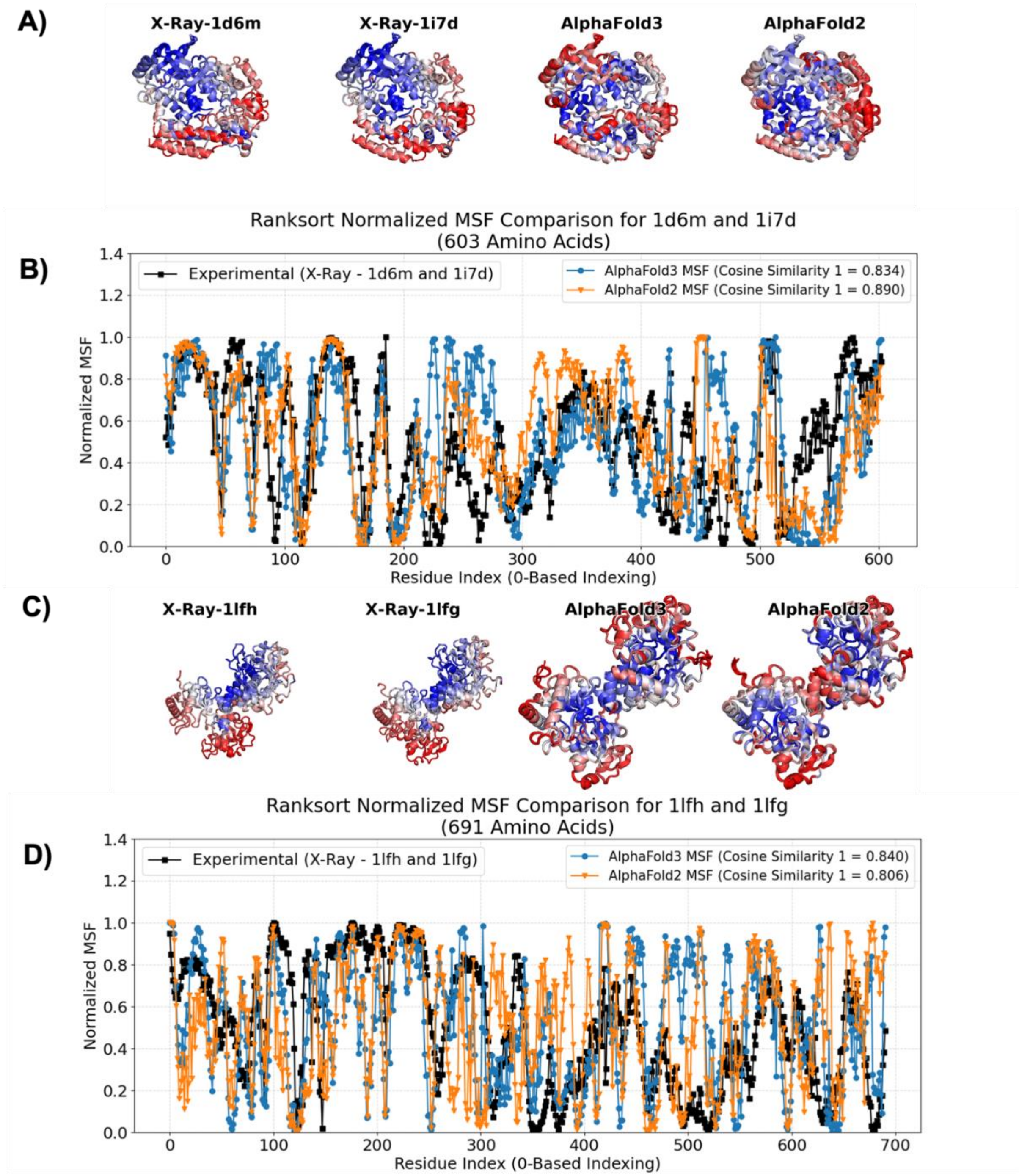


**Figure S4.** Ranksorted mean squared fluctuations (MSF) from dual conformations of X-Ray structures and three deep learning structure prediction methods (AlphaFold3 and AlphaFold2) for two proteins with more than 500 amino acids. A) Projections of the ranksorted MSF onto protein structures for 1d6m-1i7d. Blue-White-Red color palette is used for the projections, where blue indicates low flexibility and red indicates high flexibility. B) 2D comparison of the experimental and the computed MSF for 1d6m-1i7d protein pair. Black line (with squares) is for the experimental data, blue line (with circles) is for AlphaFold3 and orange line (with inverse triangles) is for AlphaFold2. C) Projections of the ranksorted MSF onto protein structures for 1lfh-1lfg. D) 2D comparison of the experimental and the computed MSF for 1lfh-1lfg protein pair. Black line (with squares) is for the experimental data, blue line (with circles) is for AlphaFold3 and orange line (with inverse triangles) is for AlphaFold2.

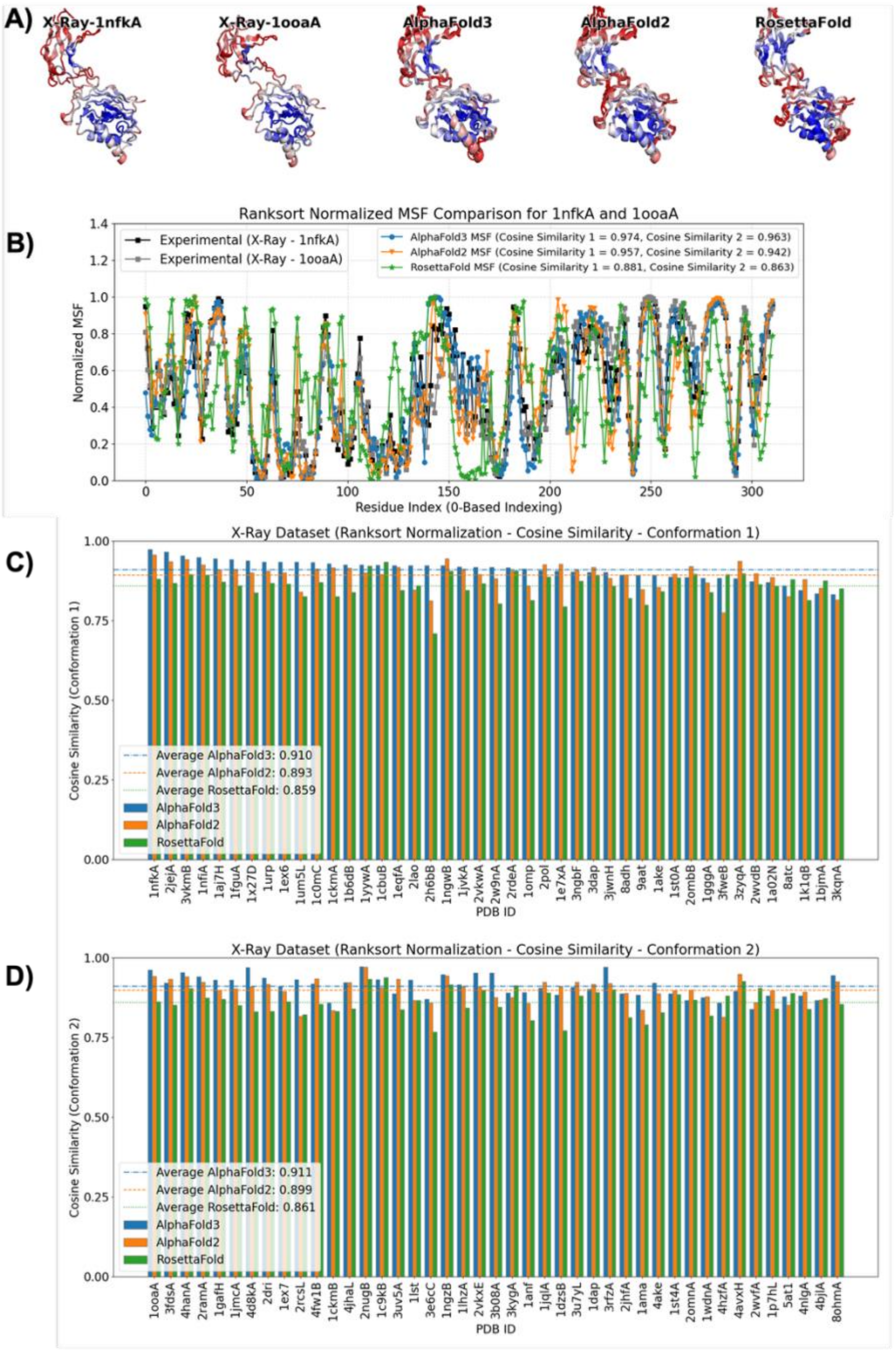


**Figure S5.** Ranksorted mean squared fluctuations (MSF) from <u>all normal modes of X-Ray structures</u> and three deep learning structure prediction methods (AlphaFold3, AlphaFold2 and RosettaFold). A) Projections of the ranksorted MSF onto protein structures for 1nfk chain A and 1ooa chain A. Blue-White-Red color palette is used for the projections, where blue indicates low flexibility and red indicates high flexibility. B) 2D comparison of the experimental and the computed MSF for 1nfk chain A and 1ooa chain A. Black line (with squares) is for the experimental data, blue line (with circles) is for AlphaFold3, orange line (with inverse triangles) is for AlphaFold2 and green line (with stars) is for RosettaFold. C) Cosine similarities of the experimental and the computed MSF for the first conformation of 43 proteins in the X-Ray dataset. AlphaFold3 bars are blue, AlphaFold2 bars are orange and RosettaFold bars are green. Averages of the cosine similarities over the entire dataset are also provided as horizontal lines for AlphaFold3 (blue dot-dashed line), AlphaFold2 (orange dashed line) and RosettaFold (green dotted line). D) Cosine similarities of the experimental and the computed MSF for the second conformation of 43 proteins in the X-Ray dataset. The colors are the same as in C.

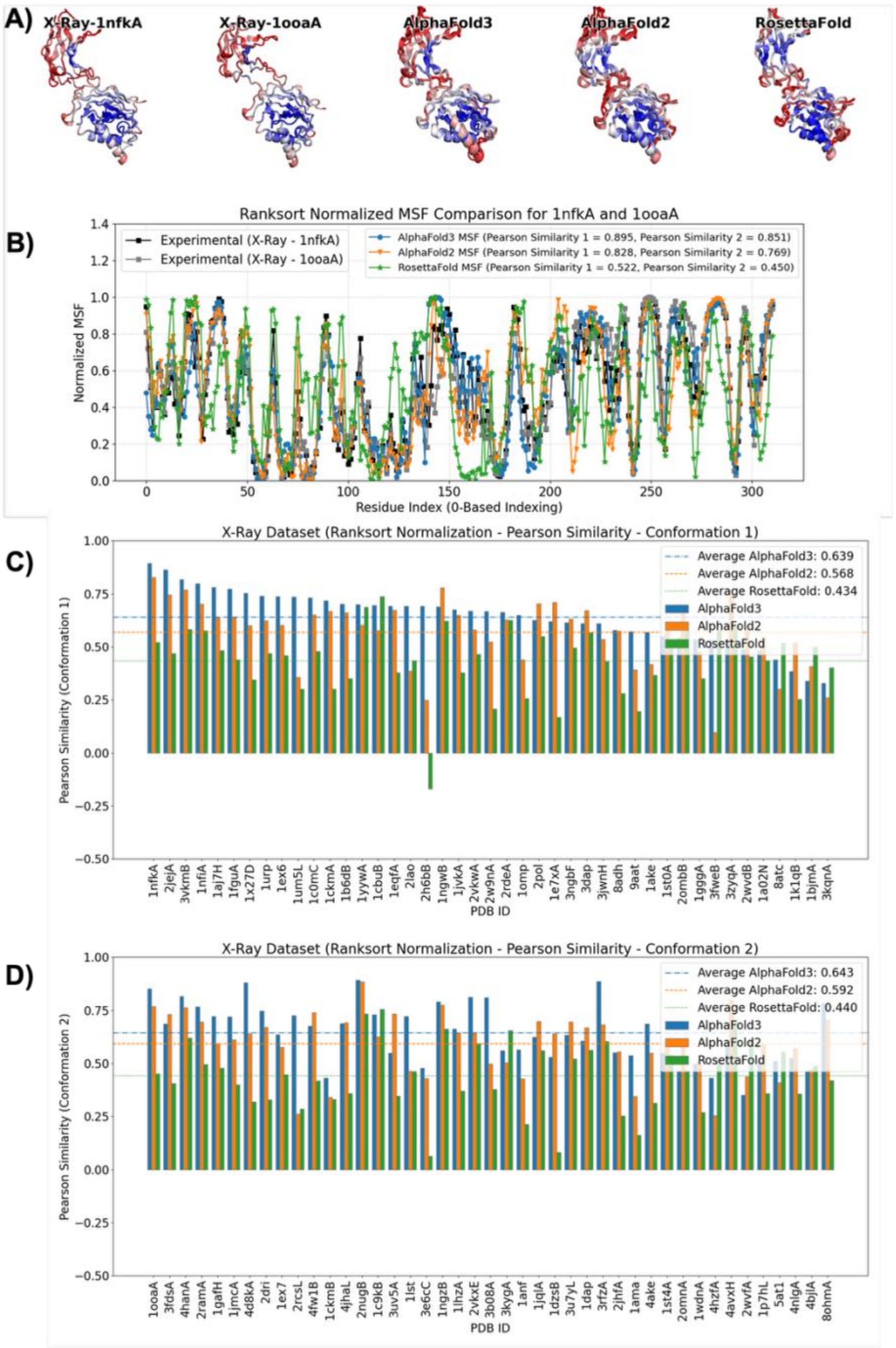


**Figure S6.** Ranksorted mean squared fluctuations (MSF) from all normal modes of X-Ray structures and three deep learning structure prediction methods (AlphaFold3, AlphaFold2 and RosettaFold). A) Projections of the ranksorted MSF onto protein structures for 1nfk chain A and 1ooa chain A. Blue-White-Red color palette is used for the projections, where blue indicates low flexibility and red indicates high flexibility. B) 2D comparison of the experimental and the computed MSF for 1nfk chain A and 1ooa chain A. Black line (with squares) is for the experimental data, blue line (with circles) is for AlphaFold3, orange line (with inverse triangles) is for AlphaFold2 and green line (with stars) is for RosettaFold. C) Pearson similarities of the experimental and the computed MSF for the first conformation of 43 proteins in the X-Ray dataset. AlphaFold3 bars are blue, AlphaFold2 bars are orange and RosettaFold bars are green. Averages of Pearson similarities over the entire dataset are also provided as horizontal lines for AlphaFold3 (blue dot-dashed line), AlphaFold2 (orange dashed line) and RosettaFold (green dotted line). D) Pearson similarities of the experimental and the computed MSF for the second conformation of 43 proteins in the X-Ray dataset. The colors are the same as in C.

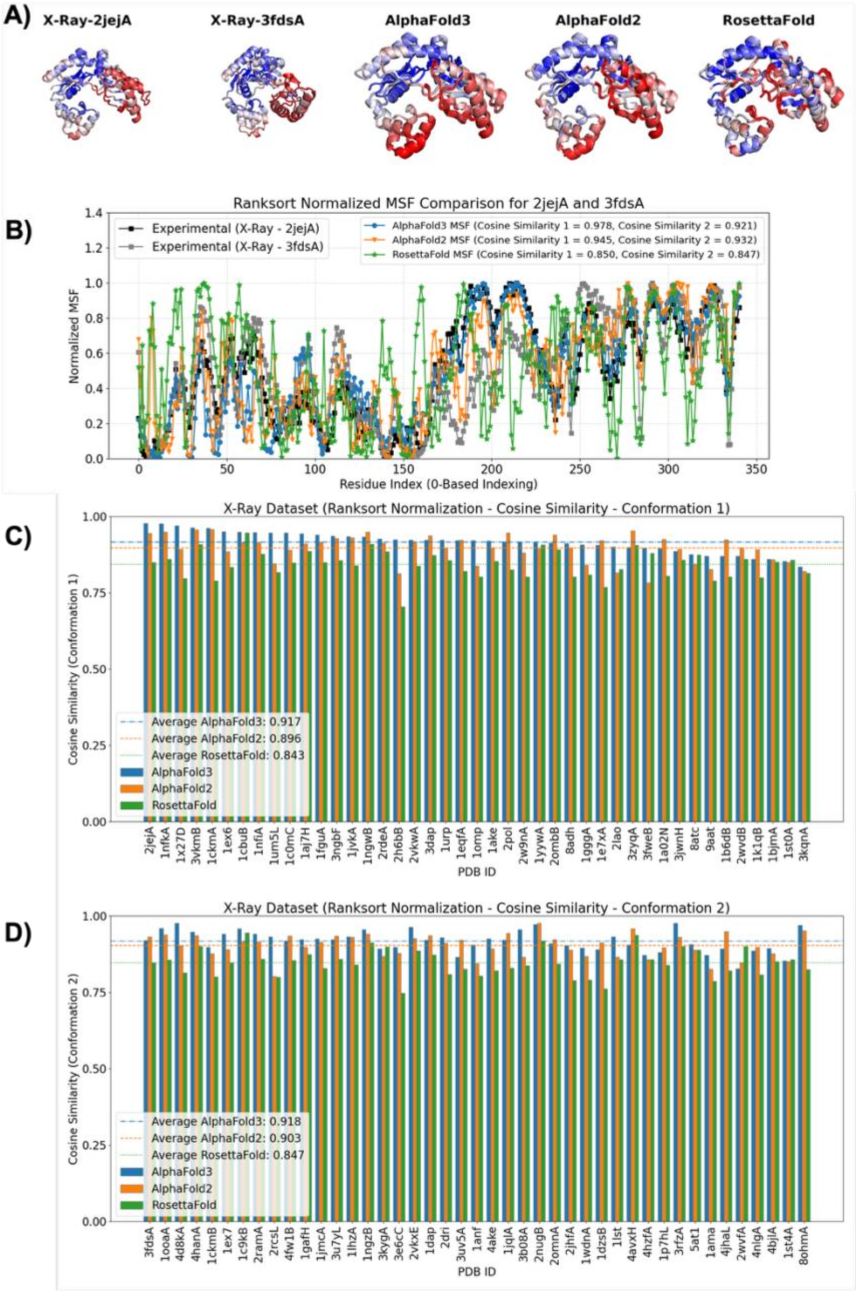


**Figure S7.** Ranksorted mean squared fluctuations (MSF) from the lowest frequency 10 normal modes of X-Ray structures and three deep learning structure prediction methods (AlphaFold3, AlphaFold2 and RosettaFold). A) Projections of the ranksorted MSF onto protein structures for 2jej chain A and 3fds chain A. Blue-White-Red color palette is used for the projections, where blue indicates low flexibility and red indicates high flexibility. B) 2D comparison of the experimental and the computed MSF for 2jej chain A and 3fds chain A. Black line (with squares) is for the experimental data, blue line (with circles) is for AlphaFold3, orange line (with inverse triangles) is for AlphaFold2 and green line (with stars) is for RosettaFold. C) Cosine similarities of the experimental and the computed MSF for the first conformation of 43 proteins in the X-Ray dataset. AlphaFold3 bars are blue, AlphaFold2 bars are orange and RosettaFold bars are green. Averages of the cosine similarities over the entire dataset are also provided as horizontal lines for AlphaFold3 (blue dot-dashed line), AlphaFold2 (orange dashed line) and RosettaFold (green dotted line). D) Cosine similarities of the experimental and the computed MSF for the second conformation of 43 proteins in the X-Ray dataset. The colors are the same as in C.

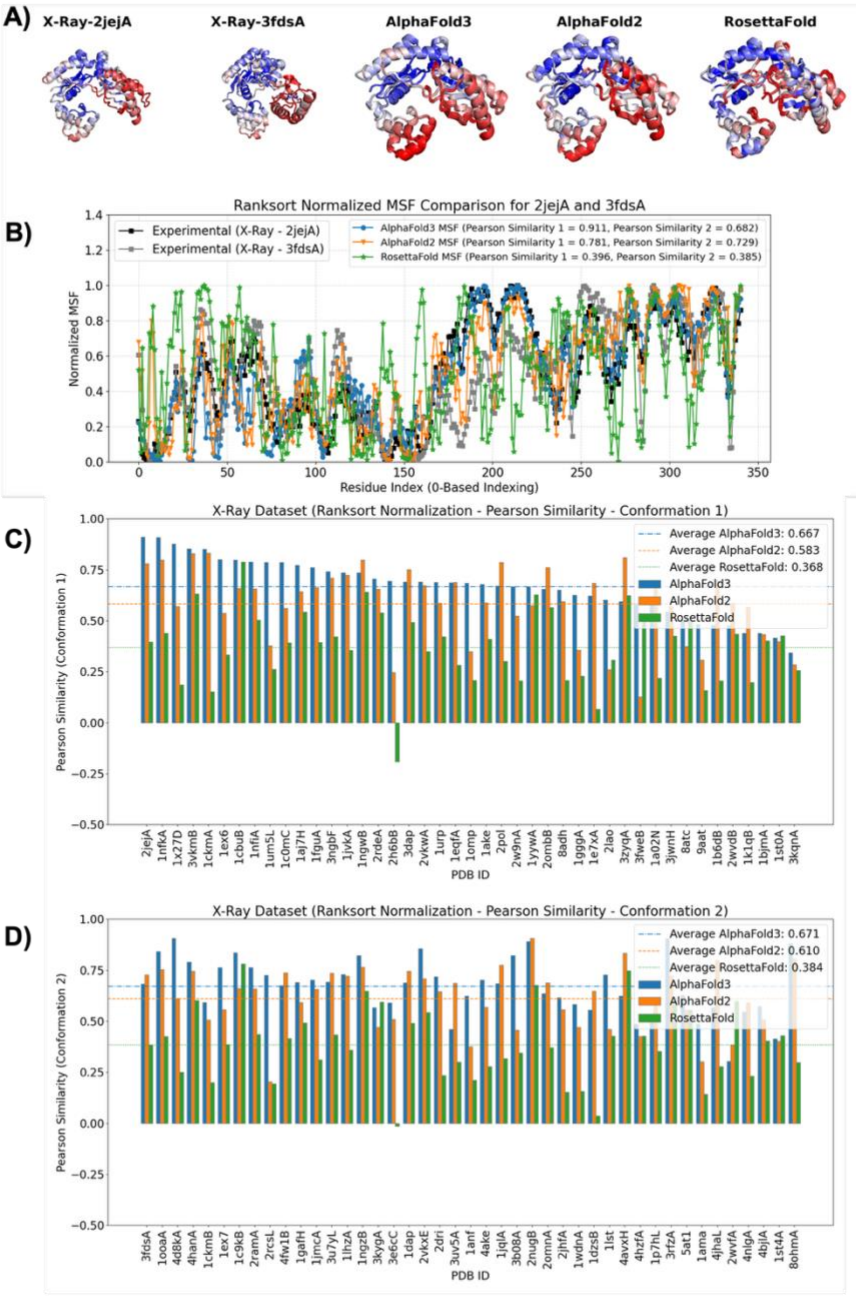


**Figure S8.** Ranksorted mean squared fluctuations (MSF) from <u>the lowest frequency 10 normal modes of X-Ray structures</u> and three deep learning structure prediction methods (AlphaFold3, AlphaFold2 and RosettaFold). A) Projections of the ranksorted MSF onto protein structures for 2jej chain A and 3fds chain A. Blue-White-Red color palette is used for the projections, where blue indicates low flexibility and red indicates high flexibility. B) 2D comparison of the experimental and the computed MSF for 2jej chain A and 3fds chain A. Black line (with squares) is for the experimental data, blue line (with circles) is for AlphaFold3, orange line (with inverse triangles) is for AlphaFold2 and green line (with stars) is for RosettaFold. C) Pearson similarities of the experimental and the computed MSF for the first conformation of 43 proteins in the X-Ray dataset. AlphaFold3 bars are blue, AlphaFold2 bars are orange and RosettaFold bars are green. Averages of Pearson similarities over the entire dataset are also provided as horizontal lines for AlphaFold3 (blue dot-dashed line), AlphaFold2 (orange dashed line) and RosettaFold (green dotted line). D) Pearson similarities of the experimental and the computed MSF for the second conformation of 43 proteins in the X-Ray dataset. The colors are the same as in C.

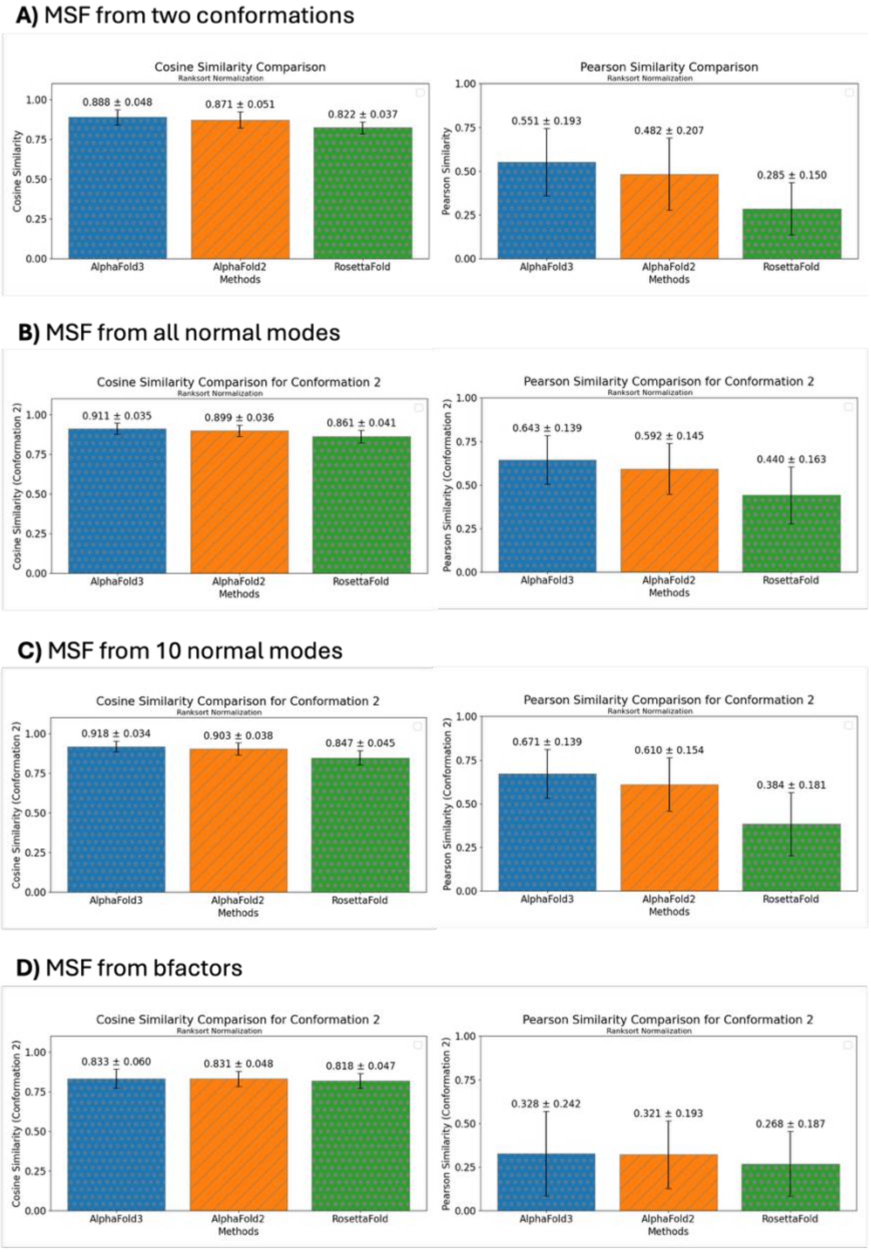


**Figure S9.** Averages of the similarities for different approaches to obtain MSF from the second conformation set of the X-Ray experimental data. The average cosine similarities are given in the left panel and the average Pearson similarities are provided in the right panel. The average was taken over the similarities of 43 proteins. Blue bars with gray circles are for AlphaFold3, orange bars with gray stripes are for AlphaFold2 and green bars with gray stars are for RosettaFold. A) Two experimental protein conformations were used for the MSF calculations. B) All normal modes of the second conformations was used for the MSF calculations. Only Calpha atoms were used for normal mode analysis. C) Only 10 lowest eigenvalue normal modes of each conformation were used for normal mode analysis. Only Calpha atoms were used for normal mode analysis. D) Bfactors were ranksort normalized and they were used as a proxy for the MSF.

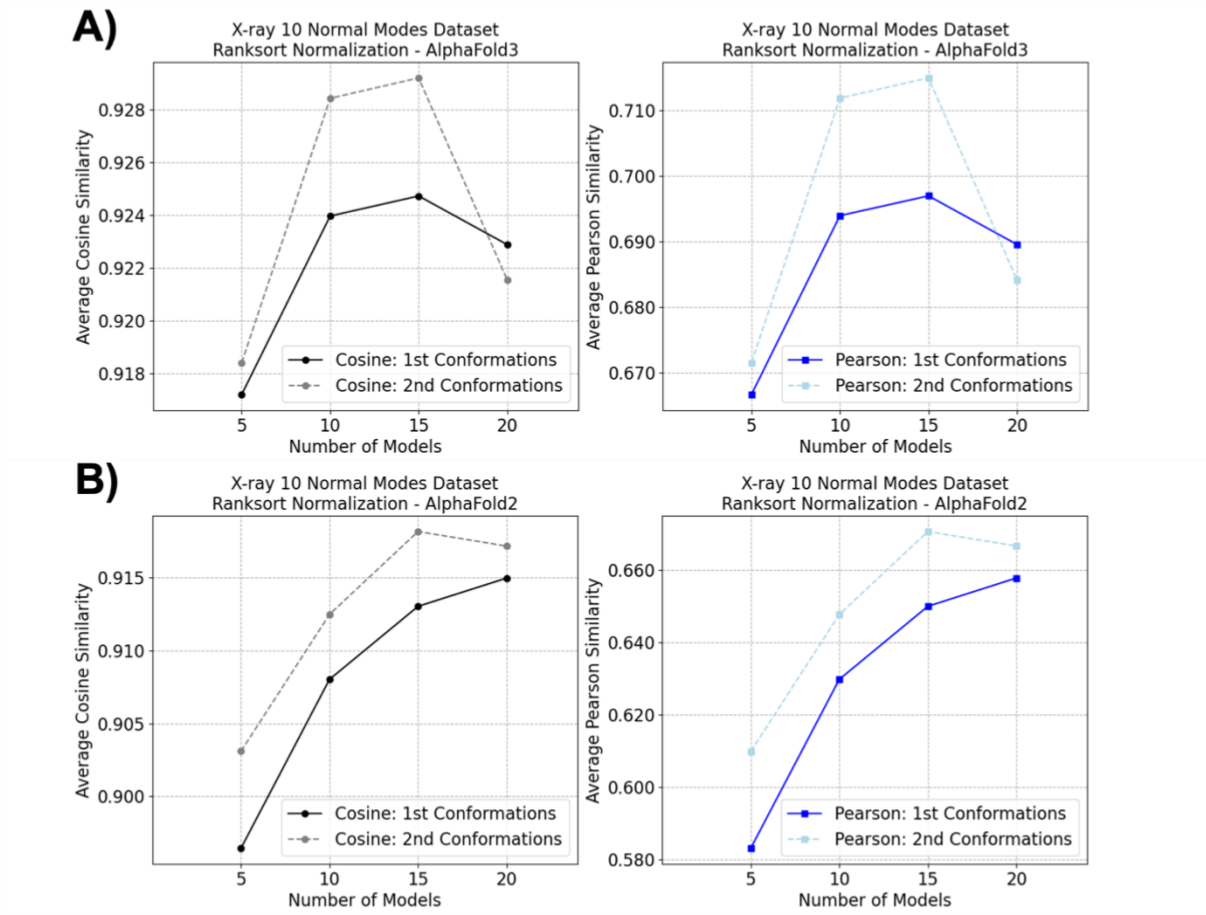


**Figure S10.** Impact of number of deep learning structural models on average similarity of X-ray 10 normal modes dataset, measured with cosine similarity (left panel, black and gray lines) and Pearson similarity (right panel, blue and light blue lines) for A) AlphaFold3 B) AlphaFold2. Continuous lines are for the first conformations set and dashed lines are for the second conformations set.

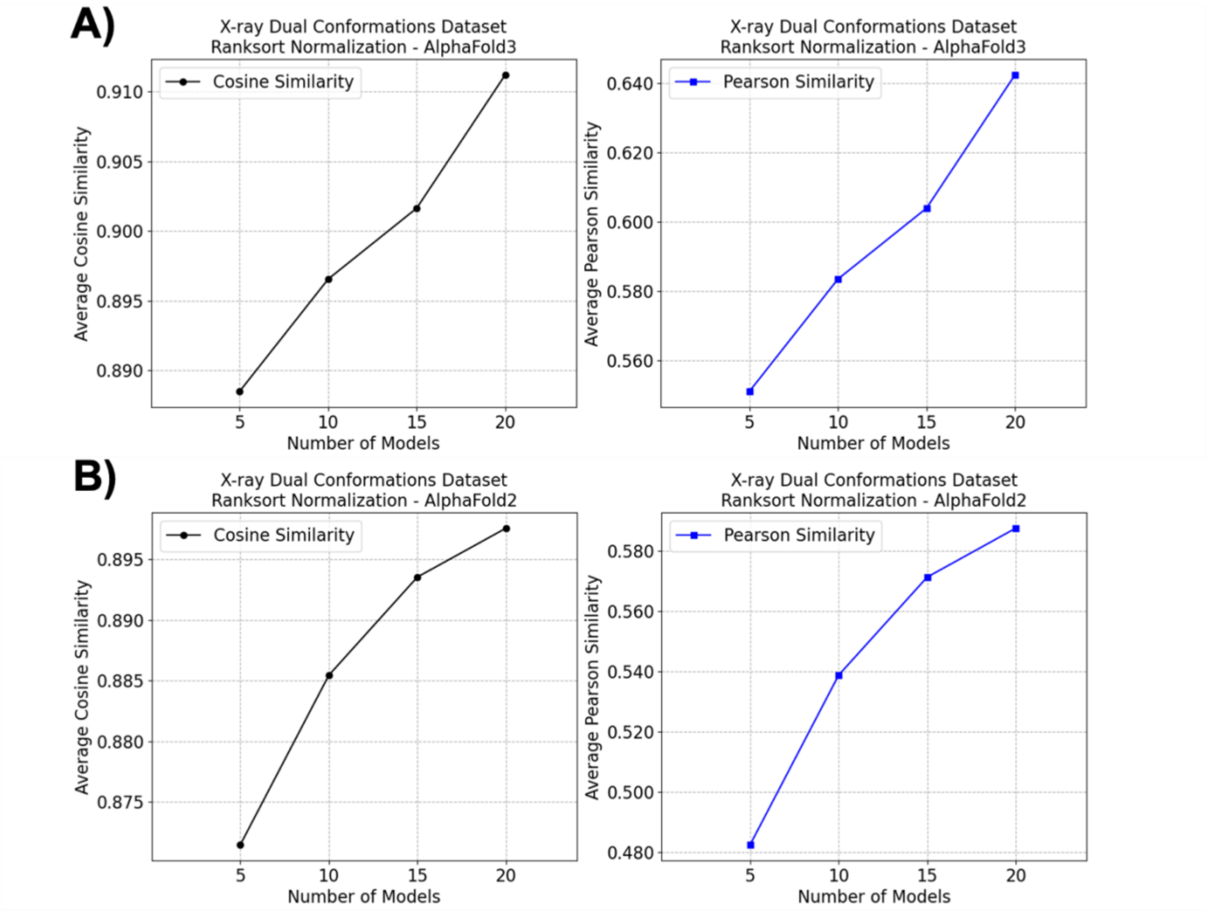


**Figure S11.** Impact of number of deep learning structural models on average similarity of X-ray dual conformations dataset, measured with cosine similarity (left panel, black line) and Pearson similarity (right panel, blue line) for A) AlphaFold3 B) AlphaFold2.

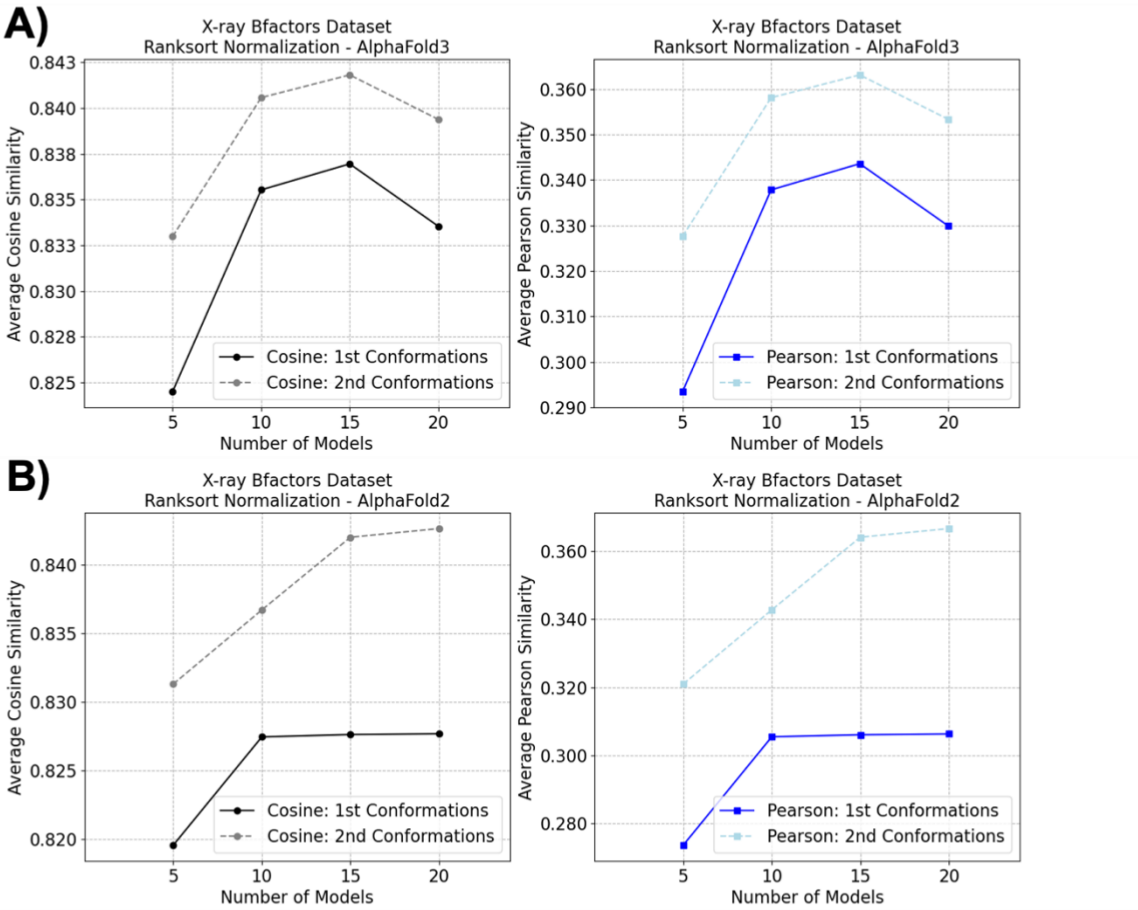


**Figure S12.** Impact of number of deep learning structural models on average similarity of X-ray bfactors dataset, measured with cosine similarity (left panel, black and gray lines) and Pearson similarity (right panel, blue and light blue lines) for A) AlphaFold3 B) AlphaFold2. Continuous lines are for the first conformations set and dashed lines are for the second conformations set.

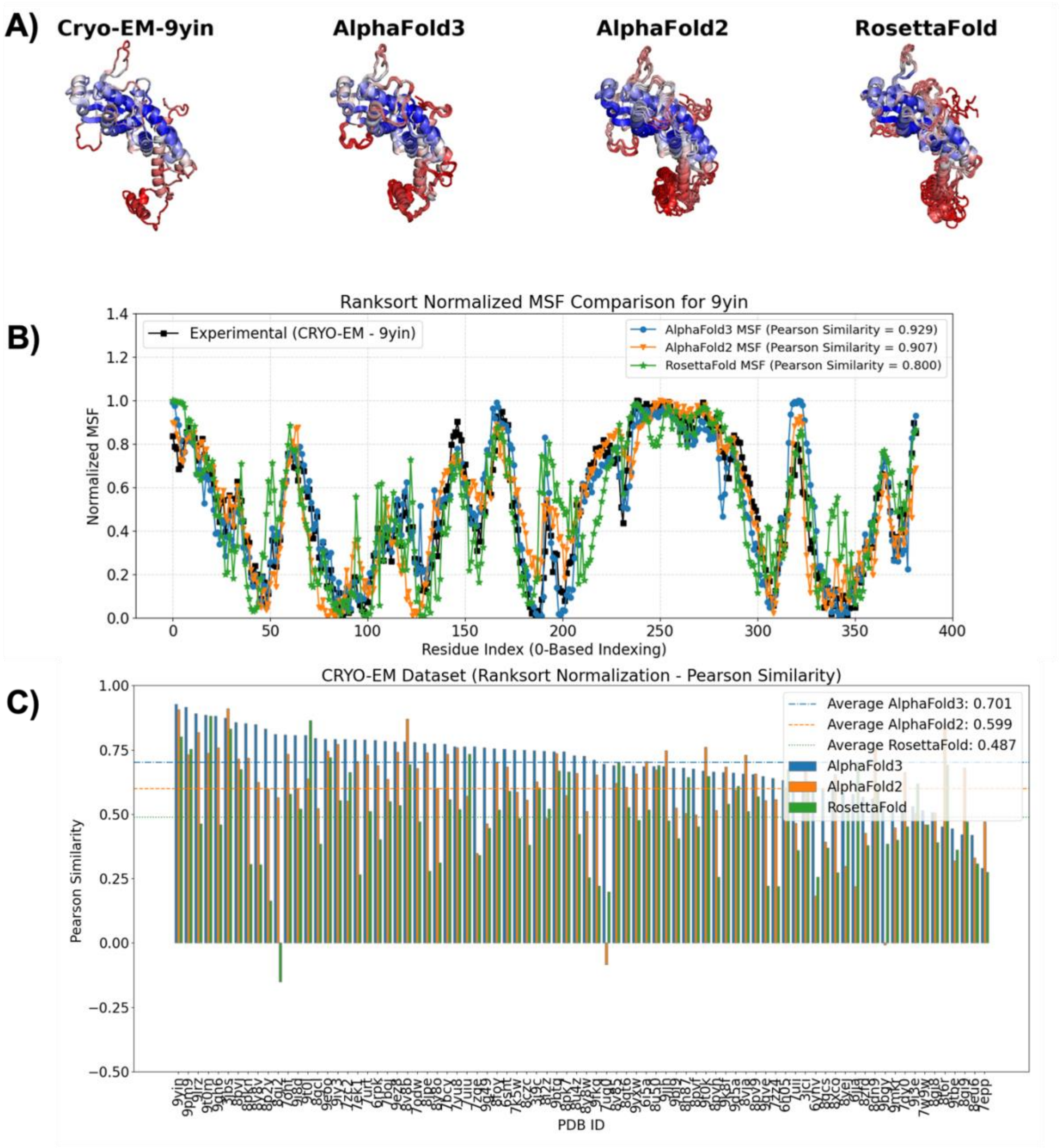


**Figure S13.** Ranksorted mean squared fluctuations (MSF) from <u>all normal modes</u> of the cryo-EM structures and three deep learning structure prediction methods (AlphaFold3, AlphaFold2 and RosettaFold). A) Projections of the ranksorted MSF onto protein structures for 9yin. Blue-White-Red color palette is used for the projections, where blue indicates low flexibility and red indicates high flexibility. B) 2D comparison of the experimental and the computed MSF for 9yin. Black line (with squares) is for the experimental data, blue line (with circles) is for AlphaFold3, orange line (with inverse triangles) is for AlphaFold2 and green line (with stars) is for RosettaFold. C) Pearson similarity of the experimental and the computed MSF for 82 proteins in the cryo-EM dataset. AlphaFold3 bars are blue, AlphaFold2 bars are orange and RosettaFold bars are green. Averages of the Pearson similarities over the entire dataset are also provided as horizontal lines for AlphaFold3 (blue dot-dashed line), AlphaFold2 (orange dashed line) and RosettaFold (green dotted line).

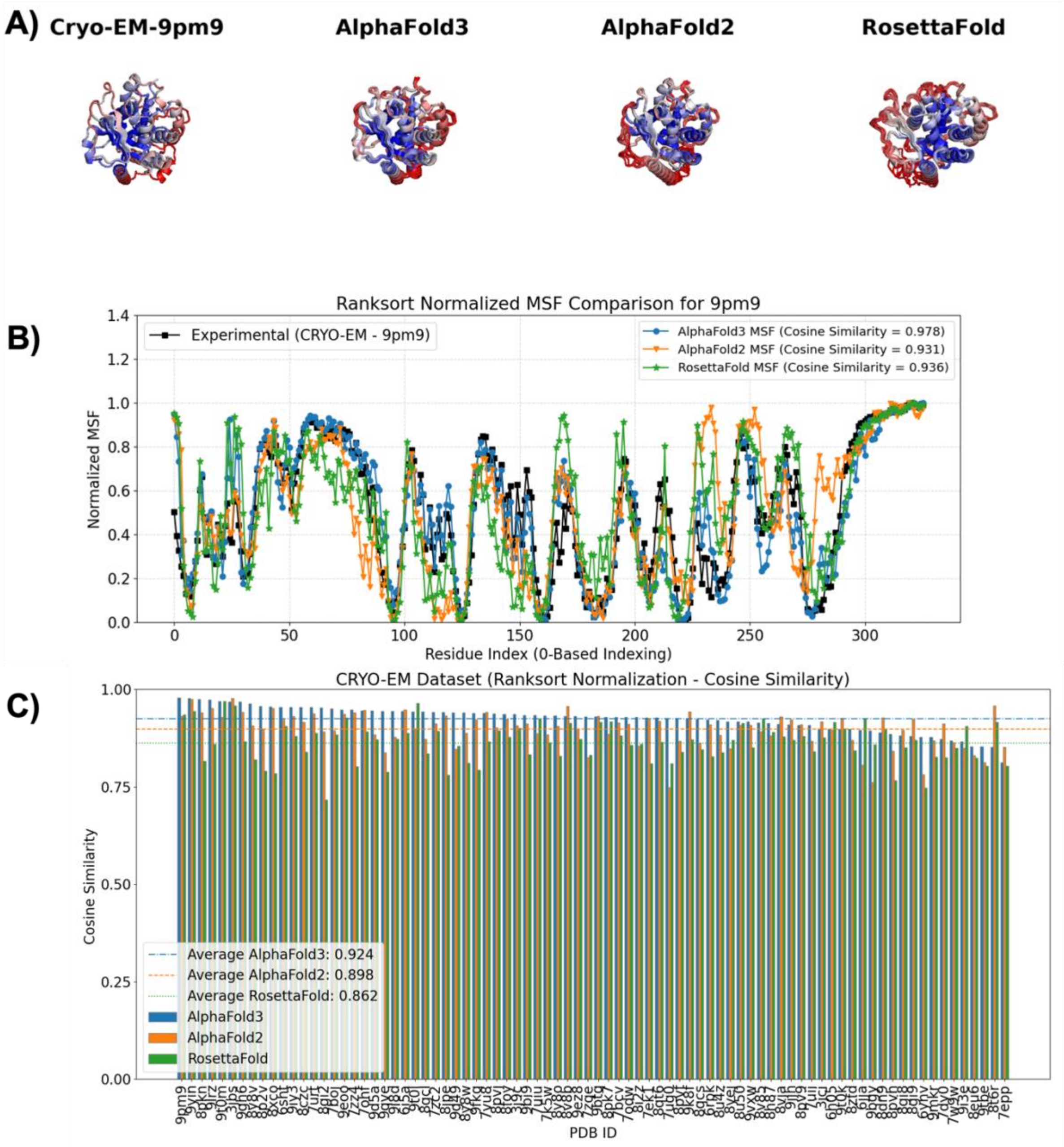


**Figure S14.** Ranksorted mean squared fluctuations (MSF) from the first 10 normal modes of the cryo-EM structures and three deep learning structure prediction methods (AlphaFold3, AlphaFold2 and RosettaFold). A) Projections of the ranksorted MSF onto protein structures for 9pm9. Blue-White-Red color palette is used for the projections, where blue indicates low flexibility and red indicates high flexibility. B) 2D comparison of the experimental and the computed MSF for 9pm9. Black line (with squares) is for the experimental data, blue line (with circles) is for AlphaFold3, orange line (with inverse triangles) is for AlphaFold2 and green line (with stars) is for RosettaFold. C) Cosine similarity of the experimental and the computed MSF for 82 proteins in the cryo-EM dataset. AlphaFold3 bars are blue, AlphaFold2 bars are orange and RosettaFold bars are green. Averages of the cosine similarities over the entire dataset are also provided as horizontal lines for AlphaFold3 (blue dot-dashed line), AlphaFold2 (orange dashed line) and RosettaFold (green dotted line).

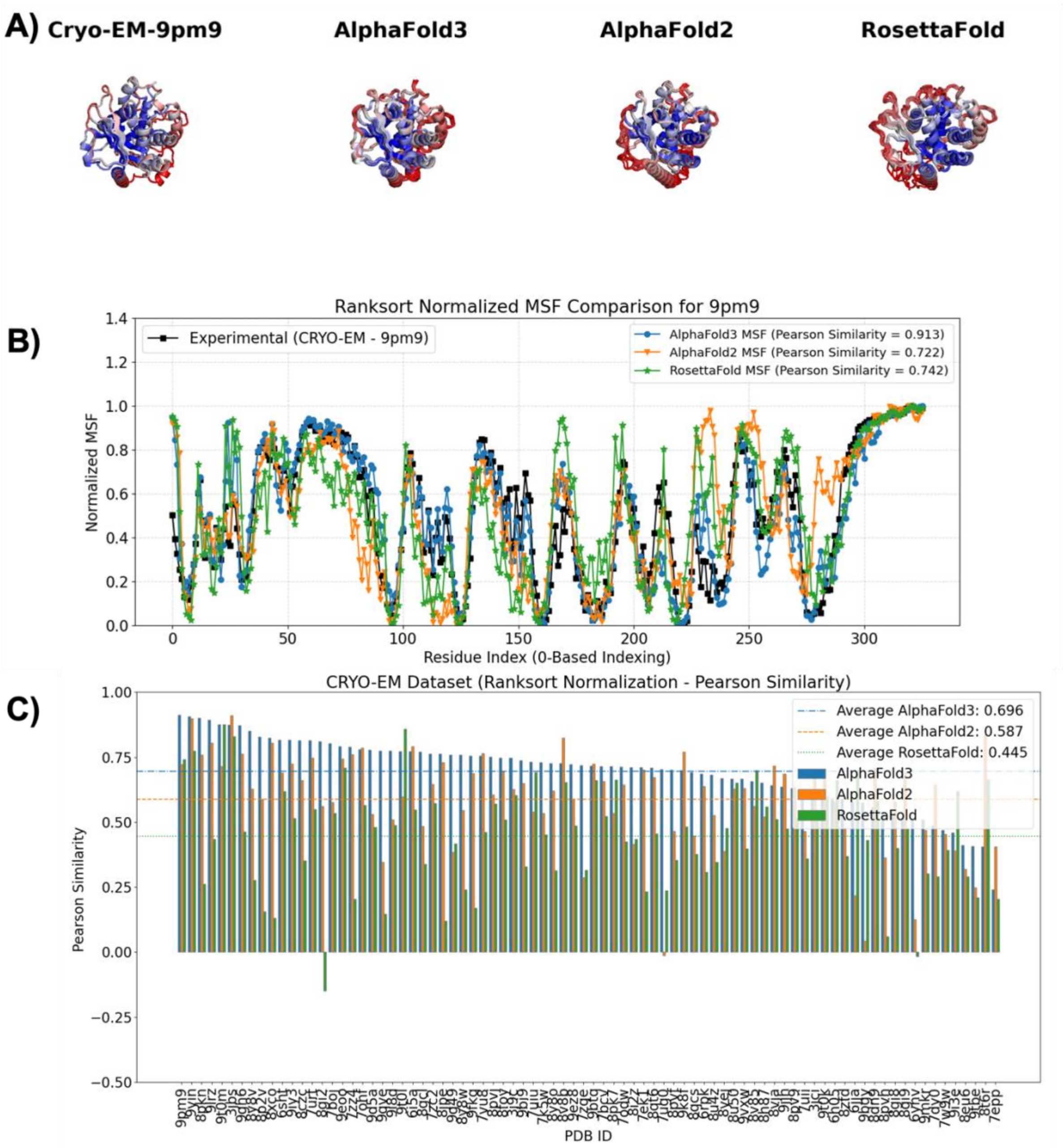


**Figure S15.** Ranksorted mean squared fluctuations (MSF) from <u>the first 10 normal modes</u> of the cryo-EM structures and three deep learning structure prediction methods (AlphaFold3, AlphaFold2 and RosettaFold). A) Projections of the ranksorted MSF onto protein structures for 9pm9. Blue-White-Red color palette is used for the projections, where blue indicates low flexibility and red indicates high flexibility. B) 2D comparison of the experimental and the computed MSF for 9pm9. Black line (with squares) is for the experimental data, blue line (with circles) is for AlphaFold3, orange line (with inverse triangles) is for AlphaFold2 and green line (with stars) is for RosettaFold. C) Pearson similarity of the experimental and the computed MSF for 82 proteins in the cryo-EM dataset. AlphaFold3 bars are blue, AlphaFold2 bars are orange and RosettaFold bars are green. Averages of the Pearson similarities over the entire dataset are also provided as horizontal lines for AlphaFold3 (blue dot-dashed line), AlphaFold2 (orange dashed line) and RosettaFold (green dotted line).

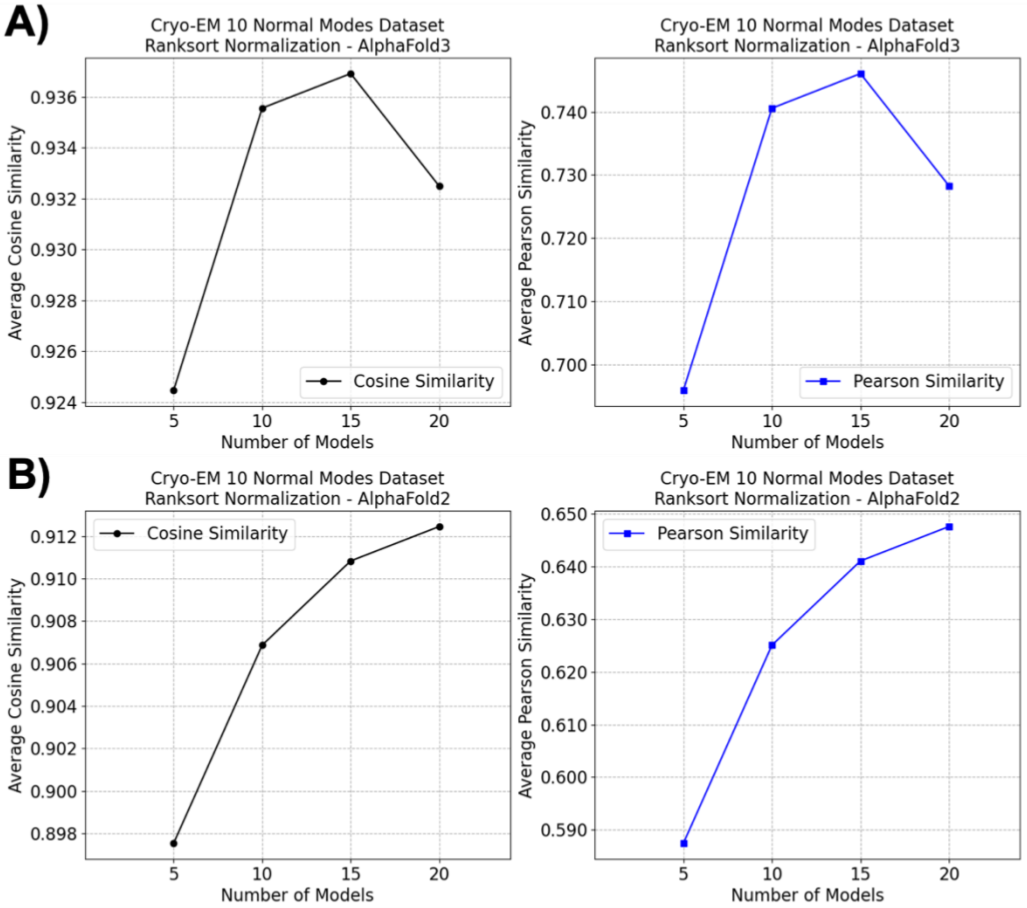


**Figure S16.** Impact of number of deep learning structural models on the cryo-EM 10 normal modes dataset, measured with cosine similarity (left panel) and Pearson similarity (right panel) for A) AlphaFold3 B) AlphaFold2.

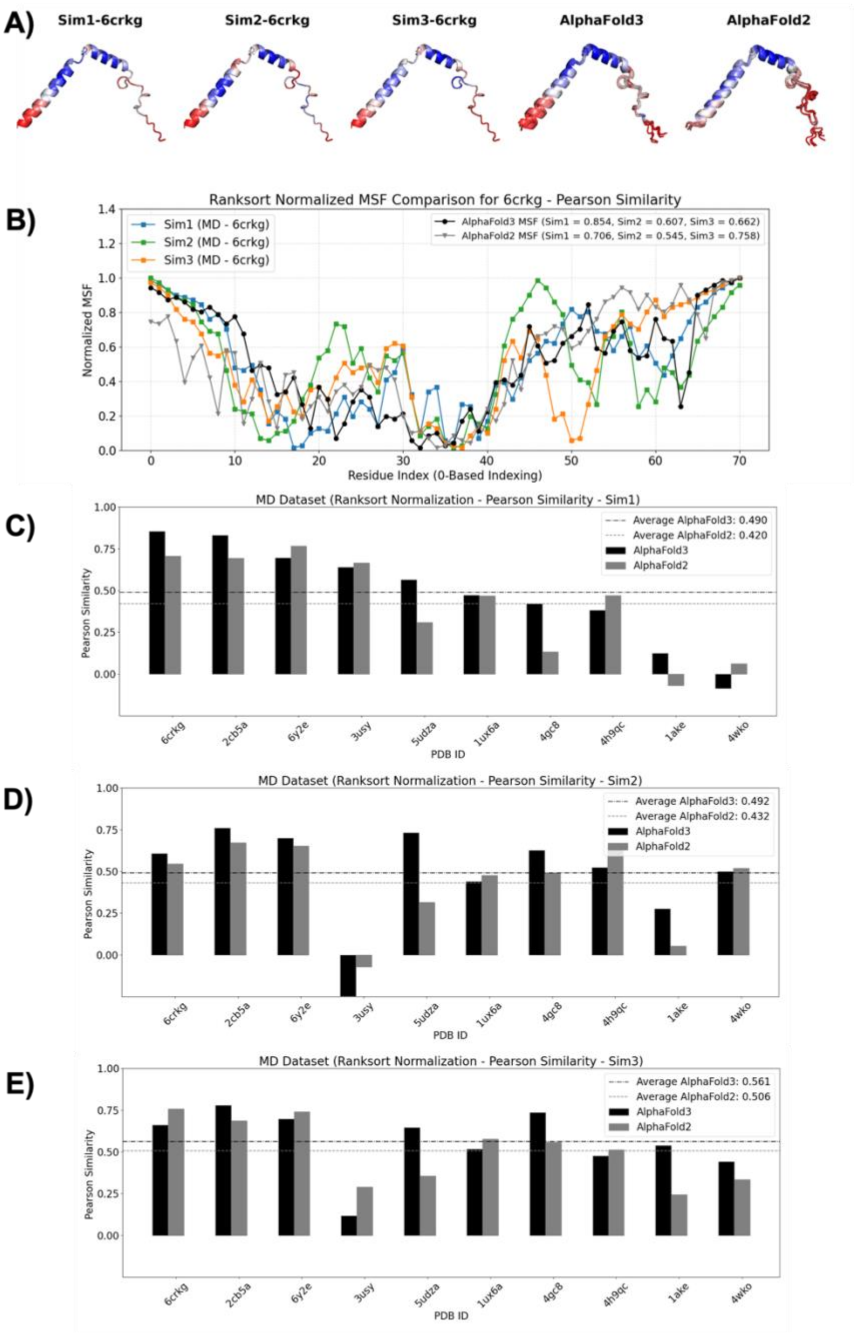


**Figure S17.** Ranksorted mean squared fluctuations (MSF) from molecular dynamics (MD) simulations and three deep learning structure prediction methods (AlphaFold3 and AlphaFold2). A) Projections of the ranksorted MSF onto protein structures for 6crk chain G. Blue-White-Red color palette is used for the projections, where blue indicates low flexibility and red indicates high flexibility. B) 2D comparison of the MD and the deep learning ensemble MSF for 6crk chain G. Blue (simulation 1), green (simulation 2) and orange (simulation 3) lines (with squares) are for the MD simulation data, black line (with circles) is for AlphaFold3, gray line (with inverse triangles) is for AlphaFold2. C) Pearson similarities of the first set of MD simulations and the deep learning ensemble MSF of 10 proteins in the MD dataset. D) Pearson similarities of the second set of MD simulations and the deep learning ensemble MSF of 10 proteins in the MD dataset. E) Pearson similarities of the third set of MD simulations and the deep learning ensemble MSF 10 proteins in the MD dataset. AlphaFold3 bars are black, AlphaFold2 bars are gray. Averages of the cosine similarities over the entire dataset are also provided as horizontal lines for AlphaFold3 (black dot-dashed line), AlphaFold2 (gray dashed line).

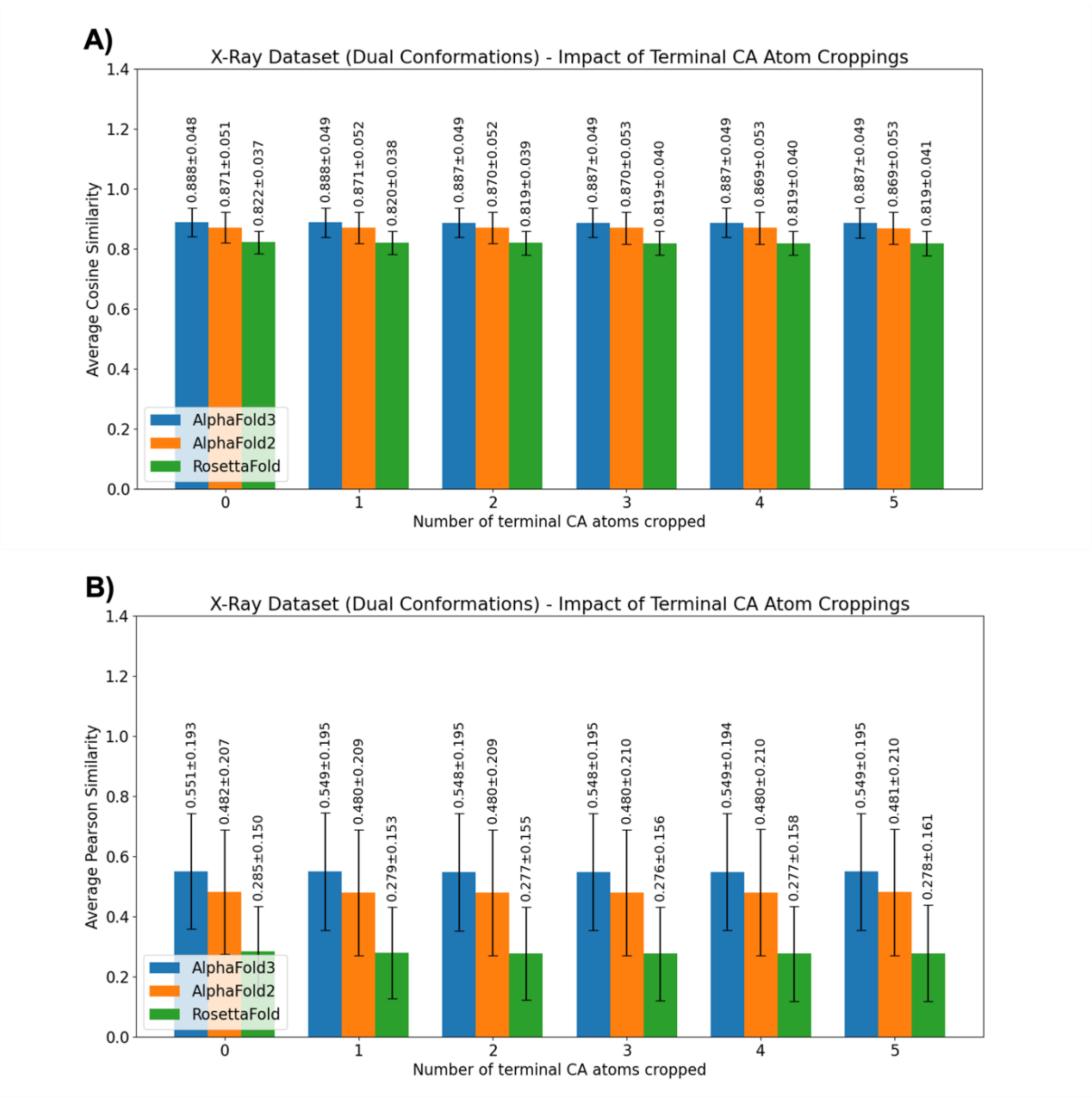


**Figure S18.** Systematic investigation of the impact of cropping MSF values of the terminal CA (Calpha) atoms for the X-Ray dual conformations dataset. A) Average cosine similarity values B) Average Pearson similarity values.

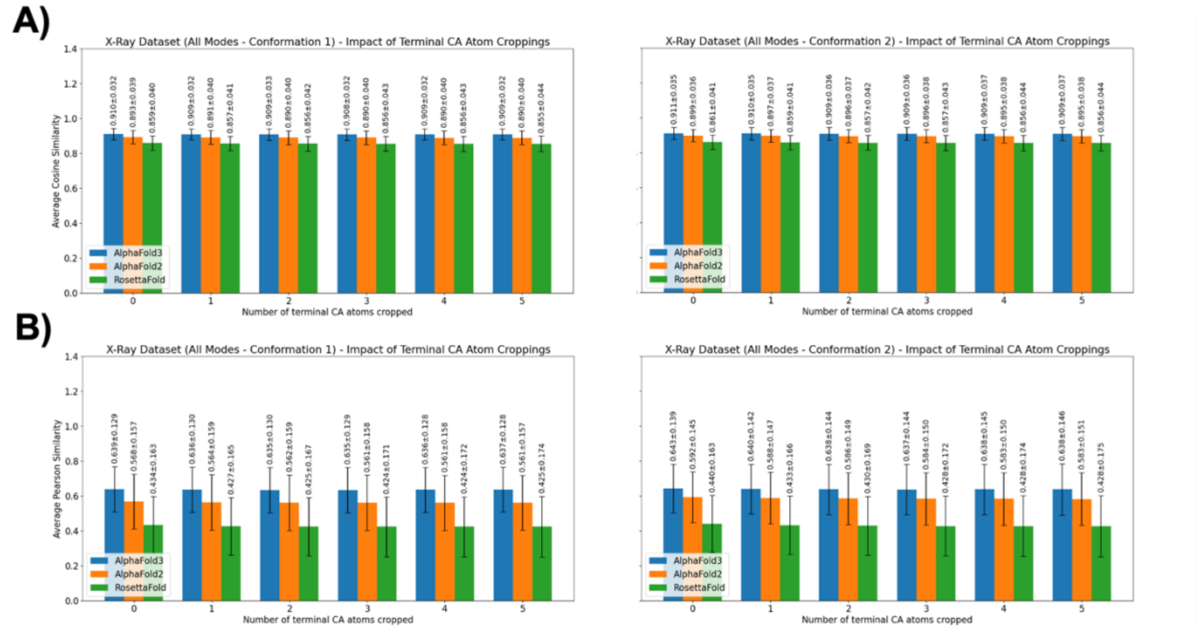


**Figure S19.** Systematic investigation of the impact of cropping MSF values of the terminal CA (Calpha) atoms for the X-Ray all normal modes dataset. The left panel is for the first conformations and the right panel is for the second conformations. A) Average cosine similarity values B) Average Pearson similarity values.

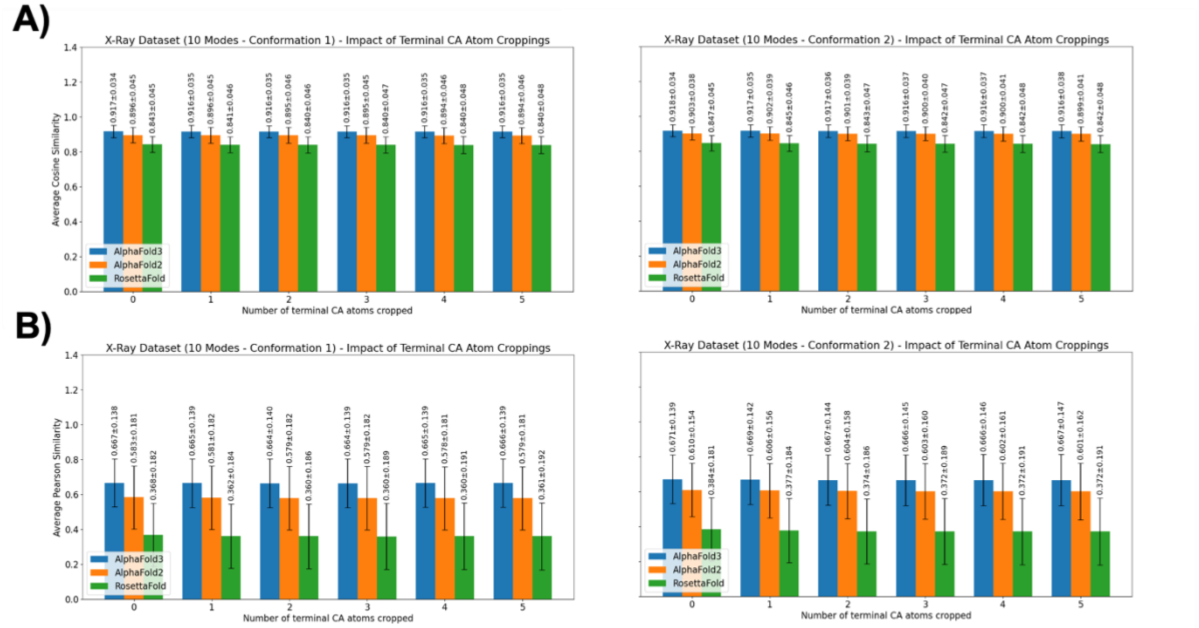


**Figure S20.** Systematic investigation of the impact of cropping MSF values of the terminal CA (Calpha) atoms for the X-Ray 10 normal modes dataset. The left panel is for the first conformations and the right panel is for the second conformations. A) Average cosine similarity values B) Average Pearson similarity values.

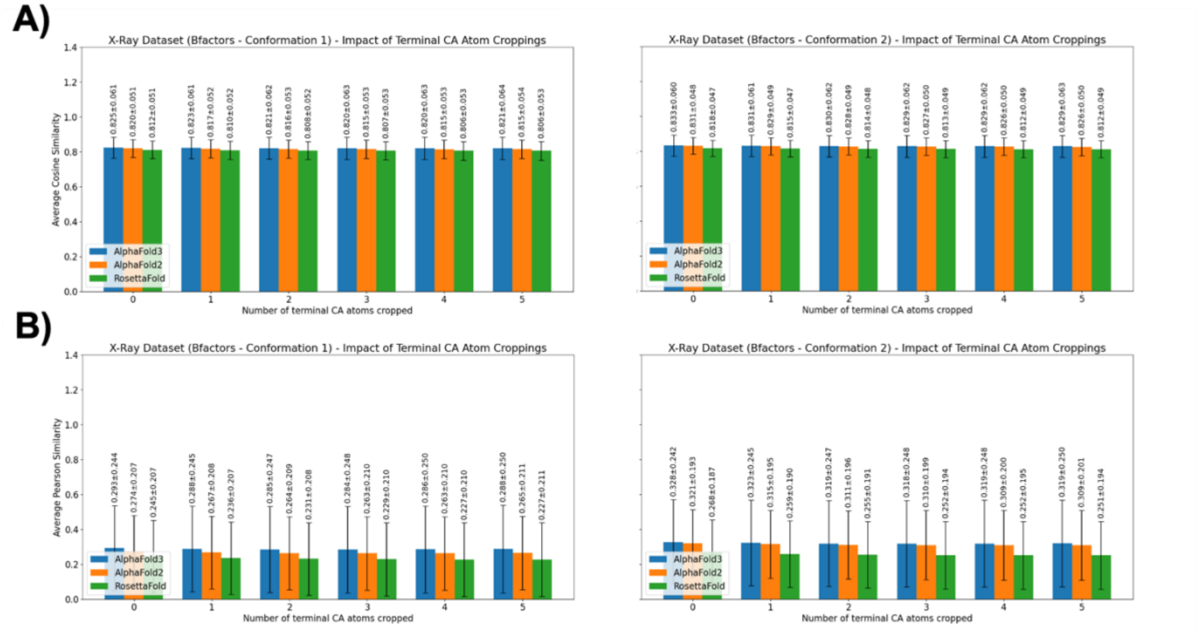


**Figure S21.** Systematic investigation of the impact of cropping MSF values of the terminal CA (Calpha) atoms for the X-Ray bfactors dataset. The left panel is for the first conformations and the right panel is for the second conformations. A) Average cosine similarity values B) Average Pearson similarity values.

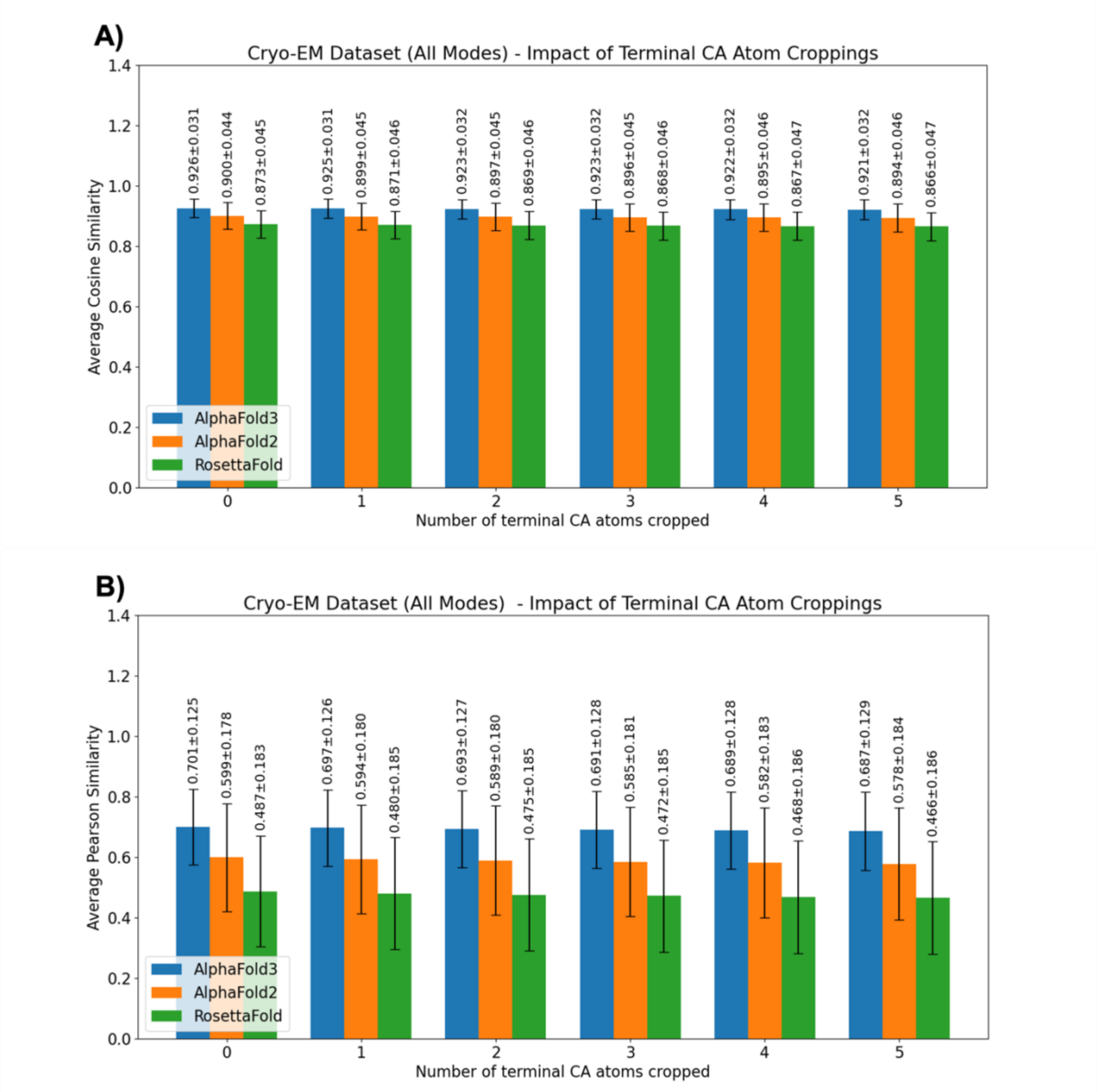


**Figure S22.** Systematic investigation of the impact of cropping MSF values of the terminal CA (Calpha) atoms for the cryo-EM all normal modes dataset. A) Average cosine similarity values B) Average Pearson similarity values.

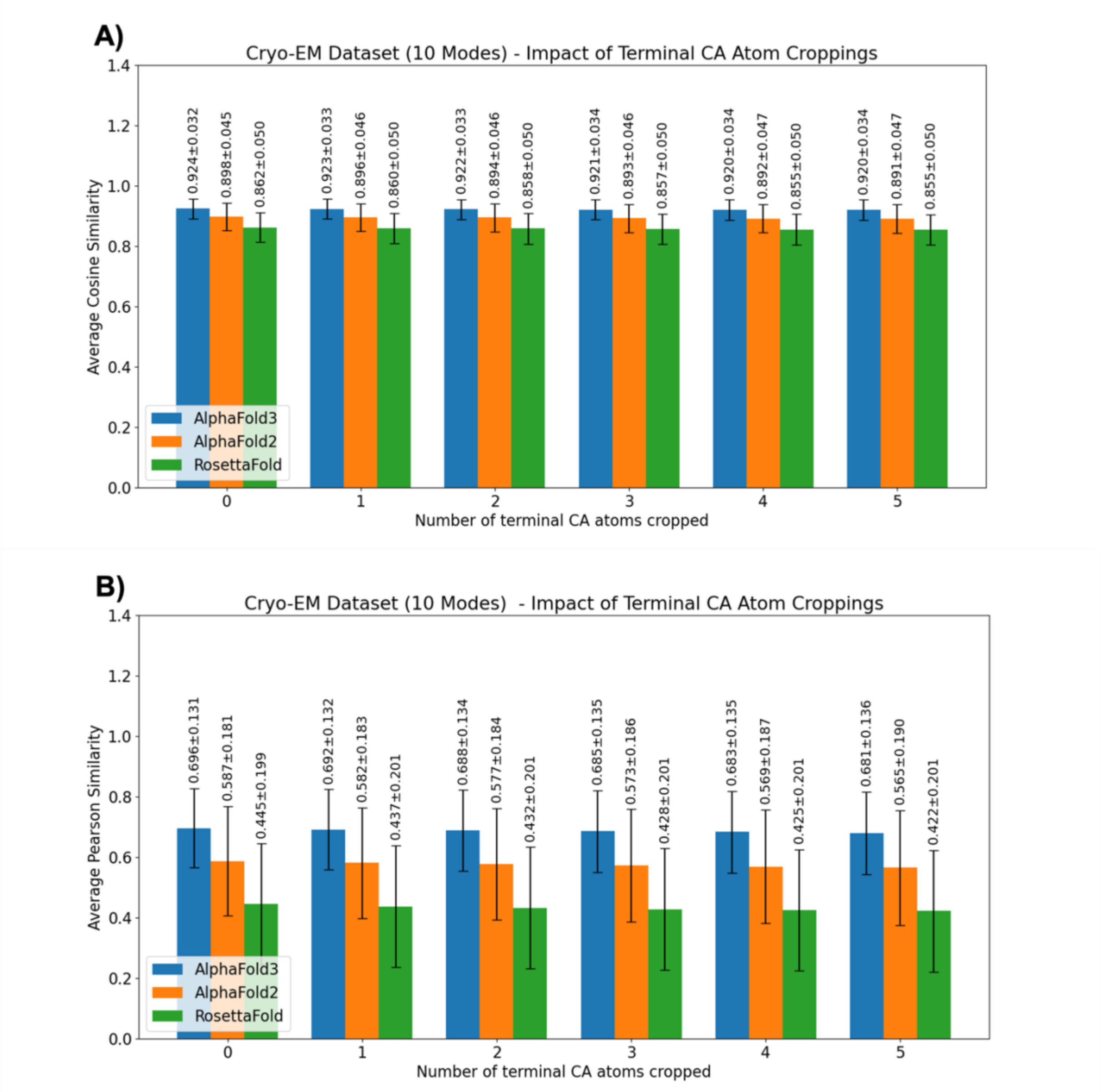


**Figure S23.** Systematic investigation of the impact of cropping MSF values of the terminal CA (Calpha) atoms for the cryo-EM 10 normal modes dataset. A) Average cosine similarity values B) Average Pearson similarity values.

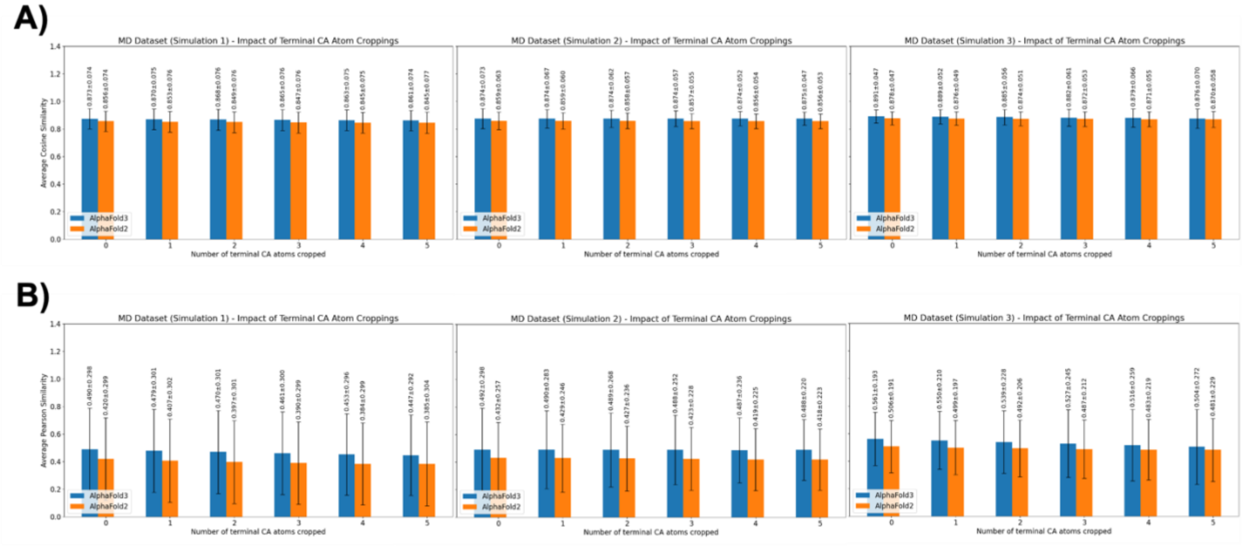


**Figure S24.** Systematic investigation of the impact of cropping MSF values of the terminal CA (Calpha) atoms for the molecular dynamics (MD) dataset. The left panel is for the first simulation set, the middle panel is for the second simulation set and the right panel is for the third simulation set. A) Average cosine similarity values B) Average Pearson similarity values.

TABLES.

**Table S1.** Protein Databank IDs, number of models and number of residues for all proteins in the NMR dataset. The proteins are ordered according to the number of residues. Three proteins highlighted with red text color have 10 models in their structures. All the remaining proteins have 20 models.

| PDB ID | Number of models | Number of residues | PDB ID | Number of models | Number of residues |
|---|---|---|---|---|---|
| **6svc** | 20 | 35 | **2kl5** | 20 | 110 |
| **6sow** | 20 | 58 | **2lta** | 20 | 110 |
| **2lx7** | 20 | 60 | **2kiw** | 20 | 111 |
| **2ma6** | 20 | 61 | **2lvb** | 20 | 112 |
| **1yez** | 10 | 68 | **2lnd** | 20 | 112 |
| **2l9r** | 20 | 69 | **1wqu** | 20 | 114 |
| **2krs** | 20 | 74 | **6gt7** | 20 | 115 |
| **2l1p** | 20 | 83 | **2kd1** | 20 | 118 |
| **2ln3** | 20 | 83 | **2ltl** | 20 | 119 |
| **2heq** | 20 | 84 | **2kvo** | 20 | 120 |
| **2kk8** | 20 | 84 | **2kcd** | 20 | 120 |
| **2kd0** | 20 | 85 | **2krt** | 20 | 121 |
| **2lml** | 20 | 87 | **2lfi** | 20 | 122 |
| **2k3d** | 20 | 87 | **2l7q** | 20 | 124 |
| **2lk2** | 20 | 89 | **2kfp** | 20 | 125 |
| **1pqx** | 10 | 91 | **2l3g** | 20 | 126 |
| **2l33** | 20 | 91 | **2l3b** | 20 | 130 |
| **2kzv** | 20 | 92 | **2lrh** | 20 | 134 |
| **2mb0** | 20 | 95 | **1vee** | 20 | 134 |
| **2kjr** | 20 | 95 | **1vdy** | 20 | 140 |
| **2m5o** | 20 | 97 | **2kkl** | 20 | 140 |
| **2lna** | 20 | 99 | **2l8v** | 20 | 143 |
| **2la6** | 20 | 99 | **2lgh** | 20 | 144 |
| **6fip** | 20 | 99 | **2k1s** | 20 | 149 |
| **2ll8** | 20 | 101 | **2m4f** | 20 | 151 |
| **2k0m** | 20 | 104 | **2jxp** | 20 | 155 |
| **2mql** | 20 | 105 | **2l06** | 20 | 155 |
| **2k75** | 20 | 106 | **2lah** | 20 | 160 |
| **2ltm** | 20 | 107 | **2lak** | 20 | 160 |
| **2kob** | 20 | 108 | **2l82** | 20 | 162 |
| **2khd** | 20 | 108 | **2m47** | 20 | 163 |
| **2rn7** | 20 | 108 | **2k3a** | 20 | 155 |
| **2lxu** | 20 | 108 | **2m7u** | 10 | 165 |
| **2kbn** | 20 | 109 | **2b3w** | 20 | 168 |
| **2mk2** | 20 | 109 | **2lf2** | 20 | 175 |

**Table S2.** Protein Databank IDs of two conformations (as PDB ID1 and PDB ID2), number of residues and RMSD between the conformations for all proteins in the X-Ray dataset. The fifth uppercase character in each PDB ID denotes the relevant chain. Please note that number of residues may have been modified to precisely match the pairs for some cases.

| PDB ID1 | PDB ID2 | Number of residues | RMSD (Å) |
|---|---|---|---|
| 2jejA | 3fdsA | 341 | 16.08 |
| 1st0A | 1st4A | 300 | 0.42 |
| 1um5L | 2rcsL | 214 | 5.43 |
| 3kqnA | 8ohmA | 435 | 5.78 |
| 2vkwA | 2vkxE | 195 | 8.31 |
| 2w9nA | 3b08A | 148 | 10.26 |
| 8atc | 5at1 | 310 | 2.35 |
| 1urp | 2dri | 271 | 4.06 |
| 1eqfA | 3uv5A | 249 | 10.75 |
| 1bjmA | 4bjlA | 215 | 2.21 |
| 3dap | 1dap | 320 | 0.29 |
| 1c0mC | 4fw1B | 214 | 14.35 |
| 1b6dB | 4jhaL | 201 | 8.29 |
| 2h6bB | 3e6cC | 225 | 15.73 |
| 1cbuB | 1c9kB | 180 | 3.11 |
| 3vkmB | 4hanA | 290 | 6.17 |
| 1x27D | 4d8kA | 162 | 10.56 |
| 2pol | 1jqlA | 366 | 1.99 |
| 1yywA | 2nugB | 218 | 11.94 |
| 3jwnH | 3rfzA | 279 | 14.65 |
| 2rdeA | 3kygA | 224 | 9.44 |
| 1jvkA | 1lhzA | 213 | 1.02 |
| 8adh | 2jhfA | 374 | 1.28 |
| 1aj7H | 1gafH | 161 | 5.05 |
| 1ake | 4ake | 214 | 7.13 |
| 1ngwB | 1ngzB | 215 | 5.17 |
| 1e7xA | 1dzsB | 129 | 3.40 |
| 1ckmA | 1ckmB | 317 | 3.49 |
| 1ex6 | 1ex7 | 186 | 3.64 |
| 1gggA | 1wdnA | 220 | 5.34 |
| 9aat | 1ama | 401 | 1.66 |
| 1k1qB | 4nlgA | 306 | 6.03 |
| 2wvdB | 2wvfA | 134 | 7.77 |
| 2ombB | 2omnA | 216 | 10.35 |
| 1a02N | 1p7hL | 280 | 16.43 |
| 2lao | 1lst | 238 | 4.70 |
| 1omp | 1anf | 370 | 3.77 |
| 1nfiA | 2ramA | 272 | 10.35 |
| 1nfkA | 1ooaA | 311 | 5.35 |
| 3ngbF | 3u7yL | 207 | 5.73 |
| 3fweB | 4hzfA | 200 | 11.04 |
| 3zyqA | 4avxH | 216 | 5.02 |
| 1fguA | 1jmcA | 238 | 8.30 |

**Table S3.** Protein Databank IDs and number of residues for all proteins in the cryo-EM dataset.

| PDB ID | Number of residues | PDB ID | Number of residues |
|---|---|---|---|
| 7uiu | 85 | 8h87 | 259 |
| 7zqe | 98 | 7k5w | 264 |
| 7dy0 | 119 | 8dn9 | 264 |
| 9btq | 124 | 7boj | 265 |
| 8pk7 | 136 | 7odw | 265 |
| 8gi2 | 147 | 9i8d | 265 |
| 6i5a | 153 | 7bcv | 266 |
| 6jja | 154 | 8u50 | 267 |
| 8pv9 | 154 | 7w9w | 270 |
| 8pvj | 155 | 9tbe | 272 |
| 9ez8 | 157 | 8vja | 275 |
| 8p2v | 161 | 9g49 | 276 |
| 7epp | 167 | 9lrz | 280 |
| 9jjh | 168 | 8zfd | 294 |
| 7ohf | 169 | 8t6r | 302 |
| 7vu8 | 170 | 8u4z | 313 |
| 6sht | 174 | 8czc | 315 |
| 9i3e | 174 | 8jpe | 317 |
| 3jbs | 175 | 8eu6 | 318 |
| 7uii | 182 | 9pm9 | 326 |
| 9qve | 184 | 7ek1 | 336 |
| 9iy3 | 189 | 8pvf | 337 |
| 3jci | 190 | 9fkq | 362 |
| 8v85 | 197 | 9yin | 382 |
| 9gh6 | 200 | 9bgy | 389 |
| 6rpk | 201 | 8qt6 | 399 |
| 9d5a | 212 | 8xco | 416 |
| 9mkr | 214 | 6ynv | 417 |
| 9eoo | 215 | 8qcs | 417 |
| 7urt | 221 | 7ug0 | 418 |
| 9k8f | 228 | 3j9c | 423 |
| 6h05 | 236 | 8pkn | 423 |
| 8gcl | 239 | 7zz4 | 463 |
| 8gi8 | 240 | 7zc2 | 471 |
| 8gi9 | 240 | 8v8b | 476 |
| 8yej | 240 | 9t0l | 480 |
| 8y8v | 245 | 9t0m | 481 |
| 8y8o | 252 | 8pvh | 493 |
| 8i2z | 254 | 9bi9 | 497 |
| 9vxw | 254 | 8foy | 500 |
| 8y8w | 257 | 9t0k | 500 |

**Table S4.** Protein Databank IDs, number of residues, number of frames and simulation times for all proteins in the molecular dynamics simulations dataset. The fifth uppercase character in each PDB ID denotes the relevant chain.

| PDB ID | Number of residues | Number of Frames | Simulation Time (ns) |
|---|---|---|---|
| **1ake** | 214 | 1001 | 200 |
| **6y2e** | 306 | 1001 | 1000 |
| **3usy** | 81 | 1001 | 1000 |
| **4gc8** | 180 | 1001 | 1000 |
| **4wkO** | 230 | 1001 | 1000 |
| **6crkG** | 71 | 1001 | 100 |
| **4h9qC** | 212 | 1001 | 100 |
| **1ux6A** | 350 | 1001 | 100 |
| **2cb5A** | 453 | 1001 | 100 |
| **5udzA** | 148 | 1001 | 100 |

AUTHOR INFORMATION

**Corresponding Authors**

*Ayten Dizkirici Tekpinar,;

*Mustafa Tekpinar,

**Author Contributions**

The manuscript was written through contributions of all authors. All authors have given approval to the final version of the manuscript.

**Notes**

The authors declare no competing interests.

ACKNOWLEDGMENT

The authors thank Google Deepmind for providing AlphaFold3 weights. The numerical calculations reported in this study were partially performed using the EuroHPC Joint Undertaking (EuroHPC JU) supercomputer MareNostrum 5, hosted by the Barcelona Supercomputing Center (BSC). The computational time was provided for the project entitled "Computing Structure and Multiple Sequence Alignments of All Human Protein Isoforms with Colabfold for Proteome Scale Analyses of Human Mutations (HumanIsoVar)". Access to MareNostrum 5 was provided through a national access call coordinated by the Scientific and Technological Research Council of Turkey (TÜBİTAK). We gratefully acknowledge BSC, TÜBİTAK, and the EuroHPC JU for providing access to these resources and supporting this research. The authors used Gemini AI (v3) tool to generate only and only abstract of the text. The abstract was read and verified by the authors for scientific accuracy. The graphical abstract was generated by ChatGPT. The authors thank Prof. Ahmet YILDIRIM for critical reading of the manuscript.

### ABBREVIATIONS

ANM, Anisotropic Network Model; Cryo-EM, Cryo-Electron Microscopy; MD, Molecular Dynamics; MSA, Multiple Sequence Alignment; MSF, Mean Squared Fluctuations; NMA, Normal Mode Analysis; NMR, Nuclear Magnetic Resonance; PDB, Protein Databank; pLDDT, predicted Local Distance Difference Test; RMSD, Root Mean Squared Distance.
